# *In situ* chromatin dynamics and HIV-1 nuclear trafficking

**DOI:** 10.64898/2025.12.12.693995

**Authors:** Zhen Hou, Yao Shen, Long Chen, Juan Shen, David I. Stuart, Peijun Zhang

## Abstract

The organization and dynamics of chromatin within intact nuclei remain poorly defined, limiting understanding of how nuclear architecture influences macromolecular transport. HIV-1 provides a striking example of a large complex that must traverse this dynamic landscape to access integration sites within transcriptionally active regions. Here, we combine a functional nuclear import system with correlative cryo-electron tomography to visualize chromatin architecture and HIV-1 cores *in situ* during nuclear penetration. Native nucleosomes resolved at 5.6 Å reveal four structural classes, from compact conformations at the nuclear periphery to open, flexible forms in the interior, defining a spatially heterogeneous and dynamic chromatin landscape. Within this environment, HIV-1 cores decorated with nuclear factors exclude nearby nucleosomes and preferentially associate with open chromatin. Disruption of CPSF6–capsid interactions abolishes these associations, confining viral cores to peripheral chromatin. These findings establish an *in situ* framework linking chromatin structure and dynamics to HIV-1 nuclear trafficking and provide a structural basis for viral navigation within the dynamic nuclear landscape.

## Introduction

In human cells, the genome is organized within the nucleus as chromatin, a dynamic nucleoprotein polymer that integrates DNA packaging with gene regulation^1^. Within the confined approximately 10 µm nucleus, roughly two meters of genomic DNA are compacted through hierarchical folding of chromatin. At the molecular level, chromatin is composed of repeating arrays of nucleosomes, each consisting of ∼147 base pairs of DNA wrapped around a histone octamer containing two copies each of H2A, H2B, H3, and H4^2, 3, 4^. Linker DNA connecting adjacent nucleosomes, when bound by histone H1, forms chromatosomes that serve as the building blocks of higher-order chromatin structures. Through nucleosome-nucleosome interactions and progressive folding, chromatin establishes domains of distinct accessibility and functionally within the nucleus, balancing genome compaction with the need for molecular accessibility during various DNA metabolic processes^5, 6^. Compact, transcriptionally repressive heterochromatin is typically enriched at the nuclear periphery and around nucleoli, whereas open, transcriptionally active euchromatin predominates in the nuclear interior^7^. Interspersed among these chromatin domains are membraneless nuclear compartments, such as nuclear speckles enriched in splicing and transcriptional regulators^8, 9^. Together, chromatin and associated nuclear bodies create a densely packed yet dynamic landscape that governs genome function and constrains the movement of macromolecular assemblies within the nucleus.

Within this crowded and heterogeneous chromatin environment, HIV-1 must penetrate and navigate the nuclear interior to access the host chromatin for integration^10, 11^. To establish infection in non-dividing cells^12, 13, 14, 15^, the HIV-1 core, a conical capsid enclosing the viral genome and enzymes, must cross the nuclear envelope through the nuclear pore complex (NPC)^16, 17, 18, 19^. The HIV-1 capsid recruits host factors to mediate nuclear transport, while simultaneously supporting the completion of reverse transcription of the viral RNA genome for subsequent integration^19, 20, 21, 22, 23, 24, 25^.

Among these host factors, cleavage and polyadenylation specificity factor 6 (CPSF6) plays a pivotal role in guiding viral cores through the nuclear environment^26, 27, 28, 29, 30^. CPSF6, a component of the cleavage factor I complex involved in mRNA 3′ end processing, binds directly to the viral capsid, facilitating nuclear entry and steering viral cores away from the heterochromatin-rich nuclear periphery toward gene-dense, transcriptionally active regions within the nuclear interior^26, 27, 28, 29, 30^. Within these subnuclear compartments, reverse transcription is thought to be completed^29, 31, 32^, and viral integration is subsequently mediated by the transcriptional coactivator lens epithelium-derived growth factor p75 (LEDGF/p75)^33, 34, 35^. LEDGF binds HIV-1 integrase and tethers the pre-integration complex to nucleosomes enriched in histone H3K36me3^33, 34, 35^, thereby promoting integration within transcriptionally active chromatin domains known as speckle-associated domains (SPADs)^29, 31, 32^ and ensuring efficient proviral transcription^33, 34, 35^. Despite these insights, how viral cores physically navigate through the dynamic chromatin environment and achieve integration site targeting remains unclear.

To address this question, we combined a functional HIV-1 nuclear import system coupled with cryo-focused ion beam (cryo-FIB) milling and correlative cryo-electron tomography (cryo-ET) to directly visualize chromatin architecture and HIV-1 cores in intact T-cell nucleus during nuclear trafficking. We resolved the *in situ* structure of nucleosomes at 5.6 Å and identified four distinct classes of nucleosome conformations, with compact nucleosome conformations preferentially enriched at the nuclear periphery and open and flexible conformations predominating within the interior, defining a dynamic and heterogeneous chromatin landscape that HIV-1 must traverse. Within this nuclear environment, HIV-1 cores decorated with nuclear factors locally exclude nucleosomes within approximately 35 nm of the capsid surface, creating nucleosome-depleted zones. Surrounding these regions, nucleosomes in open conformations are enriched, whereas compact conformations are diminished, suggesting that HIV-1 traffics through locally accessible chromatin regions. Furthermore, the N74D capsid mutation abolishes the association of viral cores with nuclear factors, thereby confining them to peripheral, compact chromatin regions. Together, these findings establish a structural framework connecting chromatin dynamics to HIV-1 nuclear trafficking, laying a foundation for future studies on chromatin remodelling, viral integration, and transcription control.

## Result

### Correlative cryo-ET of HIV-1 nuclear trafficking

While the process of HIV-1 nuclear import through NPCs has been investigated^19, 23, 24, 25^, how viral cores traverse the nuclear interior and interact with host chromatin after nuclear entry remains poorly understood. To directly visualize this process, we combined our previously established HIV-1 nuclear import assay with a correlative cryo-ET imaging approach^19, 24^.

We performed the HIV-1 nuclear import assay using isolated viral cores with permeabilized CEM cells in the presence of rabbit reticulocyte lysate and an ATP-regenerating system (RRL-ATP) (Supplementary Fig. 1a) and imaged the samples by confocal fluorescent microscopy (FM). Briefly, HIV-1 cores were isolated from virions carrying near full-length genomes (Env-deficient R9 derivatives)^36^ that expressed mNeonGreen-tagged integrase (mNeonGreen-IN).

Sucrose gradient centrifugation followed by Triton X-100 “spin-through” treatment produced a discrete fluorescent band corresponding to intact cores^37^ (Supplementary Fig. 1b, inset), which was confirmed by cryo-EM imaging (Supplementary Fig. 1b). To allow cytoplasmic access while preserving nuclear integrity, CEM cells were gently permeabilized with a low concentration of digitonin (0.018%) and supplemented with RRL-ATP^38, 39^ (Supplementary Fig. 1a). Mixing the isolated HIV-1 cores with these permeabilized cells resulted in robust nuclear import, as demonstrated by confocal imaging showing abundant mNeonGreen-IN puncta within nuclei (Supplementary Fig. 1c).

We next employed a correlative cryo-FIB and cryo-ET workflow to image imported HIV-1 cores within intact nuclei^19, 24^. Cryo-FIB milling generated cell lamellae 100–150 nm thick that contained mNeonGreen-IN signals marking HIV-1 cores and SiR-DNA signals delineating nuclei. Cryo-fluorescence microscopy (cryo-FM) of the post-milled lamellae confirmed the presence of mNeonGreen-IN signals within the nucleus (Fig. 1a), guiding cryo-ET data acquisition from the corresponding regions (Fig. 1b, Supplementary Table 1-2).

**Figure 1.**
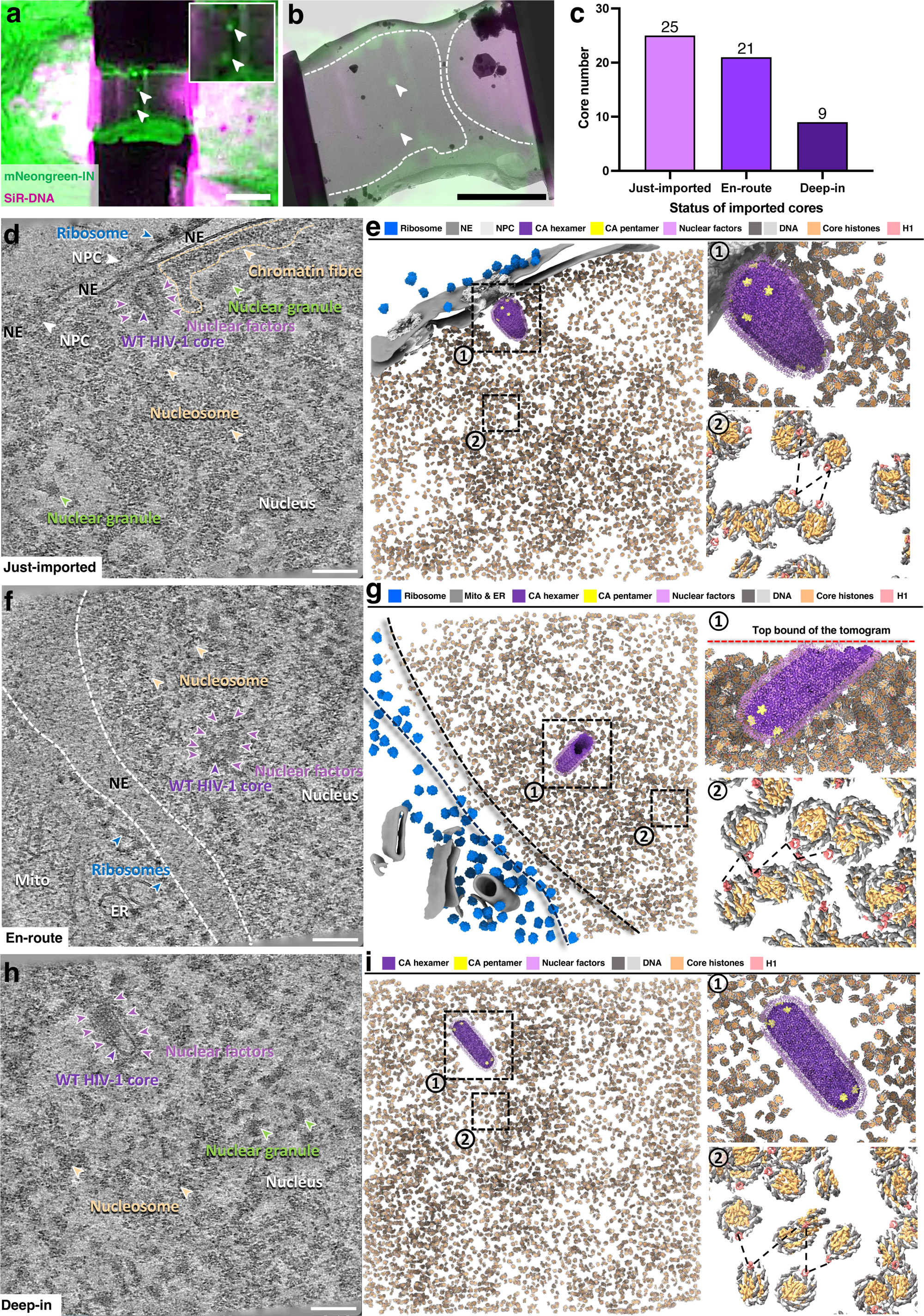
Correlative cryo-ET of nuclear trafficking of HIV-1. **a-b,** A representative cryo-fluorescence image of a lamella (**a**) and overlapped with a low magnification of cryo-EM image of the same lamella (**b**). HIV-1 cores are indicated by white arrowheads. Nuclei are outlined by white dashed lines. Inset in (**a**) is an enlarged view of the area containing HIV-1 cores. Scale bars = 5 μm. **c**, A bar chart showing the distribution of HIV-1 cores captured in nuclei at the stage of “Just-imported (<75 nm from NPC), “En-route” (between 75 nm to 1.5 µm from NPC), and “Deep-in” ( > 1.5 µm from NPC). **d**, A representative tomographic slice of a “Just-imported” cone-shaped WT HIV-1 core. The NPC, ribosomes, nucleosomes, prominent chromatin fibres, nuclear granules, and nuclear factors coating the core are labelled. The nucleus and nuclear envelope (NE) are annotated accordingly. **e**, The segmented volume of (**d**), shown as an overview (left) and zoomed-in views of the “Just-imported” WT core (upper right) and zig-zag organized nucleosomes connected by black dashed lines (right lower). HIV-1 CA hexamers and pentamers, NPC, DNA, core histones and histone H1 of nucleosomes, ribosomes, nuclear factors, and NE are segmented and mapped back with indicated colours. Of note, two subunits of the right NPC exceeds the upper bound of the tomogram, thus are not depicted. **f**, A representative tomographic slice of an “En-route” tube-shaped WT core. Labelling of the subjects is the same as in **(d)**. NE is indicated by the white dashed lines. **g**, The segmented volume of (**f**), shown as an overview (left) and zoomed-in views of the “En-route” WT core (upper right) and zig-zag organized nucleosomes connected by black dashed lines (right lower). Of note, one end of the “En-route” core exceeds the top bound (red dashed line) of the tomogram, thus is not fully depicted. HIV-1 CA hexamers and pentamers, mitochondria/ER, DNA, core histones and histone H1 of nucleosomes, ribosomes, nuclear factors are segmented and mapped back with indicated colours. NE is indicated by the black dashed lines. **h**, A representative tomographic slice of a “Deep-in” tube-shaped WT core. Labelling of the subjects is the same as in **(d)**. **i**, The segmented volume of (**h**), shown as an overview (left) and zoomed-in views of the “Deep-in” WT core (upper right) and zig-zag organized nucleosomes connected by black dashed lines (right lower). HIV-1 CA hexamers and pentamers, DNA, core histones and histone H1 of nucleosomes, and nuclear factors are mapped back and shown with indicated colours. Scale bars = 100 nm in **d, f, h**.

From 253 tomograms, we identified 55 HIV-1 cores within the nucleus (Fig. 1c and Supplementary Figure 2) exhibiting both cone- and tube-shaped morphologies (21 cone-shaped and 34 tube-shaped), consistant with our previous findings^19^. Based on their distance from the nuclear envelope, we classified the cores into three categories: just-imported (<75 nm; still associated with the NPC), en-route (75 nm–1.5 µm; detached from the NPC but within 1.5 µm of the nuclear envelope), and deep-inside (>1.5 µm from the nuclear envelope) (Fig. 1d–i and Supplementary Video 1-3). Notably, the number of detectable cores decreased with increasing depth of nuclear penetration (Fig. 1c), consistent with previous fluorescence microscopy studies of HIV-1–infected cells^29, 30^.

Nearly all nuclear HIV-1 cores displayed distinct densities on their capsid surfaces, likely corresponding to nuclear factors such as CPSF6 that mediate HIV-1 nuclear import and trafficking (Fig. 1d-i, Supplementary Fig. 2). Chromatin fibers were clearly visible beneath the nuclear envelope (Fig. 1d), where short nucleosome clutches exhibited a zigzag organization (Fig. 1d–i). Nucleosomes appeared more loosely packed in the nuclear interior (Fig. 1h) than at the periphery (Fig. 1d, f). Notably, the regions immediately surrounding HIV-1 cores were markedly depleted of nucleosomes (Fig. 1d–i, Supplementary Fig. 2), suggesting that the decorated cores may locally displace chromatin during nuclear trafficking. The mechanism underlying this chromatin exclusion during HIV-1 core transport through the nucleus remains to be elucidated.

### *In situ* structure and dynamics of nucleosomes

It is worth noting that in this HIV-1 nuclear import system, imported viral cores were relatively rare (a total of 55 cores observed across 253 tomograms, with 9 located deep within the nucleus). The majority of nucleosomes were far from HIV-1 cores and are therefore expected to remain in a largely native state. Because the HIV-1 cores were introduced exogenously for 30 minutes, the overall chromatin landscape was likely only minimally perturbed. Thus, the chromatin structures described here represent an approximately native configuration in the absence of direct association with HIV-1 cores.

Using template matching and subtomogram averaging (STA) of individual nucleosomes, we resolved four distinct nucleosome classes within the nucleus at up to 5.6 Å resolution (Supplementary Fig. 3-4). The majority of nucleosomes (Class 1, 69%) corresponded to H1-bound nucleosomes, representing canonical nucleosome core particles with histone H1 bound to linker DNA (chromatosomes). The 5.6 Å STA map revealed well-defined histone octamers with clearly resolved secondary structural features and discernible major and minor grooves of the DNA (Fig. 2a, Supplementary Fig. 3-4). This map closely fits the atomic model of *in vitro* reconstituted human nucleosomes with H1 bound on-dyad (Fig. 2b)^40^. The C-terminal tail of histone H2A was clearly resolved, extending outward from the nucleosome core to engage the linker DNA and contribute to higher-order chromatin organization^41^. Similarly, the N-terminal tails of histone H3 projected from the DNA groove, consistent with a previous high-resolution structural study^6^. These findings demonstrate the capability of *in situ* structural determination to resolve molecular details of small protein–DNA complexes (∼250 kDa) within the crowded nuclear environment.

**Figure 2.**
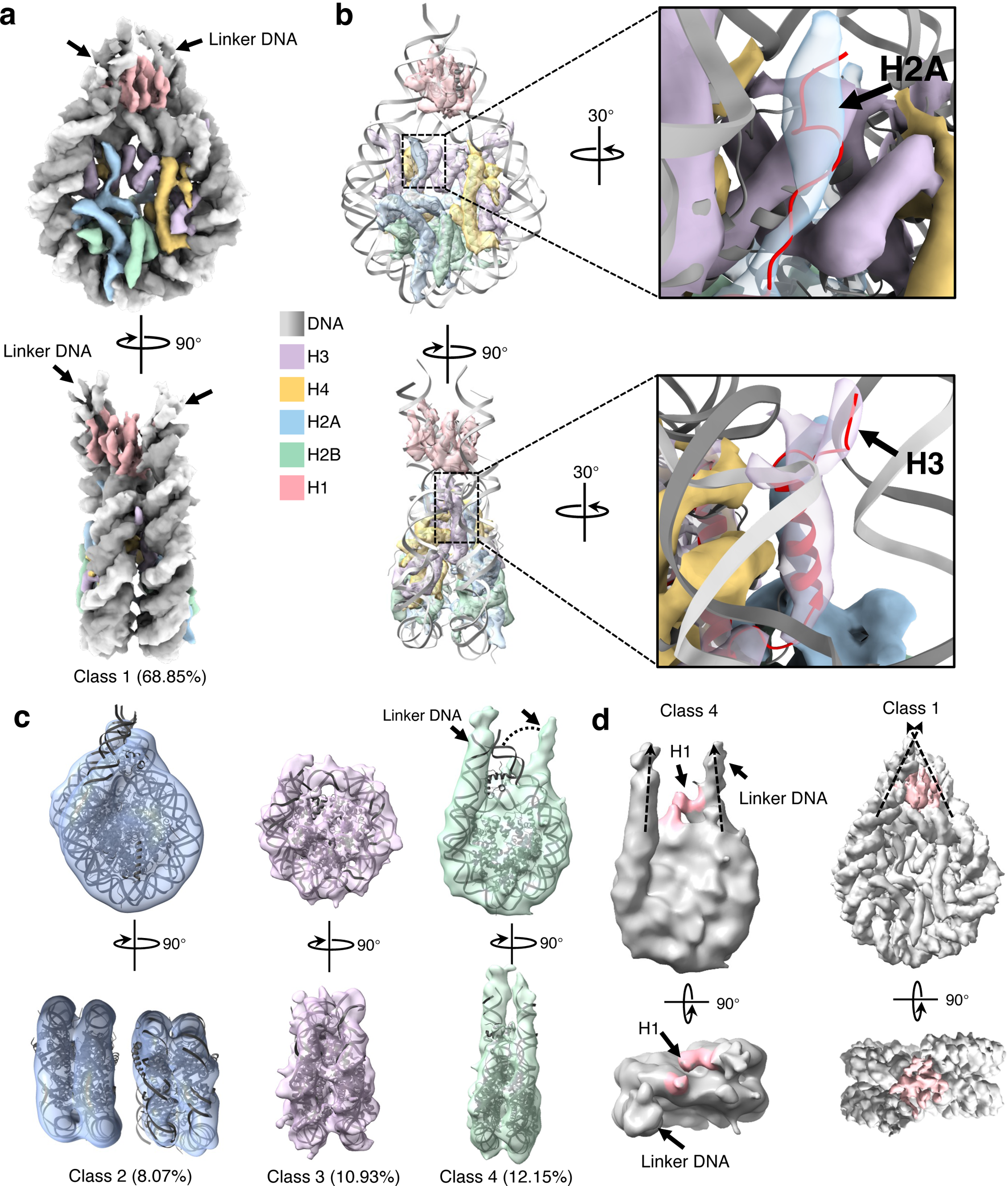
*In situ* structure and dynamics of nucleosomes. **a**, Cryo-ET STA structure of the major class (class 1) of *in situ* H1-bound nucleosome at 5.6 Å resolution. Histones and DNAs are colored as indicated. Two orthogonal views are displayed with core histones shown at 6σ, and H1 and linker DNA shown at 3σ. Linker DNA indicated by black arrows. **b**, The histone proteins are fitted with atomic model (PDB 7PEX). DNA densities are removed for clarity. Two orthogonal views are displayed with regions containing H2A and H3 tails enlarged on the right with model coloured red. **c**, Cryo-ET STA structures of class 2 (stacking H1-bound nucleosomes), class 3 (core nucleosome), and class 4 (open-linker H1-bound nucleosome). Two orthogonal views are depicted, class 2 is fitted with the atomic model (PDB 7PEX), class 3 and class 4 are fitted with the atomic model (PDB 2CV5). Class 2 is contoured at 3σ, class3 is contoured at 5σ, class 4 is contoured at 2.5σ with linker DNA indicated by black arrows. The deviation of the linker DNA density from the model is indicated by the dashed curve. **d**, Comparison between H1-bound nucleosome (class1) and open-linker H1-bound nucleosome (class 4). Two orthogonal views are depicted. The trajectory of the linker DNA is indicated by dashed arrows and H1 is coloured pink.

The second class (Class 2, 8%) revealed two parallelly stacked-nucleosomes, each associated with a histone H1 molecule (stacked H1-bound nucleosomes) (Fig. 2c, Supplementary Fig. 3-4). Focused refinement of a single nucleosome within this stack yielded a 9.9 Å map, showing that the H4 N-terminal tails extended toward adjacent nucleosomes, contacting the acidic patches on H2A–H2B dimers of neighboring nucleosomes to stabilize inter-nucleosome packing (Supplementary Fig. 5)^2, 42^. This configuration contrasts with the Class 1 H1-bound nucleosome (Supplementary Fig. 5b), where the H4 N-terminal tails fold back to form intra-nucleosomal contacts with the H3 α2 helix, as previously described^40, 43^. Notably, several histone regions at the inter-nucleosome interface, including H3 α3 and the C-terminal β3 strands of H4 and H2A, appeared disordered in the focused-refined H1-bound nucleosome within the stacked pair (Supplementary Fig. 5b). In contrast, in Class 1 nucleosomes, H4-β3 and H2A-β3 form parallel β-strands that stabilize the histone octamer and maintain nucleosome integrity^40^ (Supplementary Fig. 5b). The absence of these β-strand densities in the stacked configuration suggests that the H4-β3 tail fails to fold back, instead adopting a flexible conformation that may in turn destabilize H3-α3. Similar deviations from the folded-back H4-β3 conformation have been observed in complexes with histone chaperones during histone deposition or eviction^44, 45^.

Two additional nucleosome classes were associated with open or flexible linker DNA. Class 3 nucleosomes (core nucleosomes, 11%; 9.3 Å resolution) lacked detectable densities for H1 and linker DNA, whereas Class 4 nucleosomes (open-linker H1-bound nucleosomes, 15%; 12.1 Å resolution) featured parallel linker DNA arms, with histone H1 engaging asymmetrically with one of the linkers (Fig. 2c-d, Supplementary Figs. 3-4). Although linker histone H1 is known to stabilize linker DNA, the weaker density observed for the H1-bound linker in this class likely reflects partial H1 occupancy and conformational heterogeneity under native nuclear conditions (Fig. 2d, Supplementary Fig. 4d), reinforcing the role of H1 in regulating chromatin compaction and dynamic remodeling^6, 46, 47^.

### Chromatin architecture and nucleosome arrangement in the nucleus

To investigate how nucleosome organization contributes to higher-order chromatin architecture, we mapped nucleosome particles back into their original tomograms based on their determined positions and orientations. Neighboring nucleosomes were then connected by linker DNA segments of varying lengths (Fig. 3a), enabling analysis of their spatial relationships and overall distributions.

**Figure 3.**
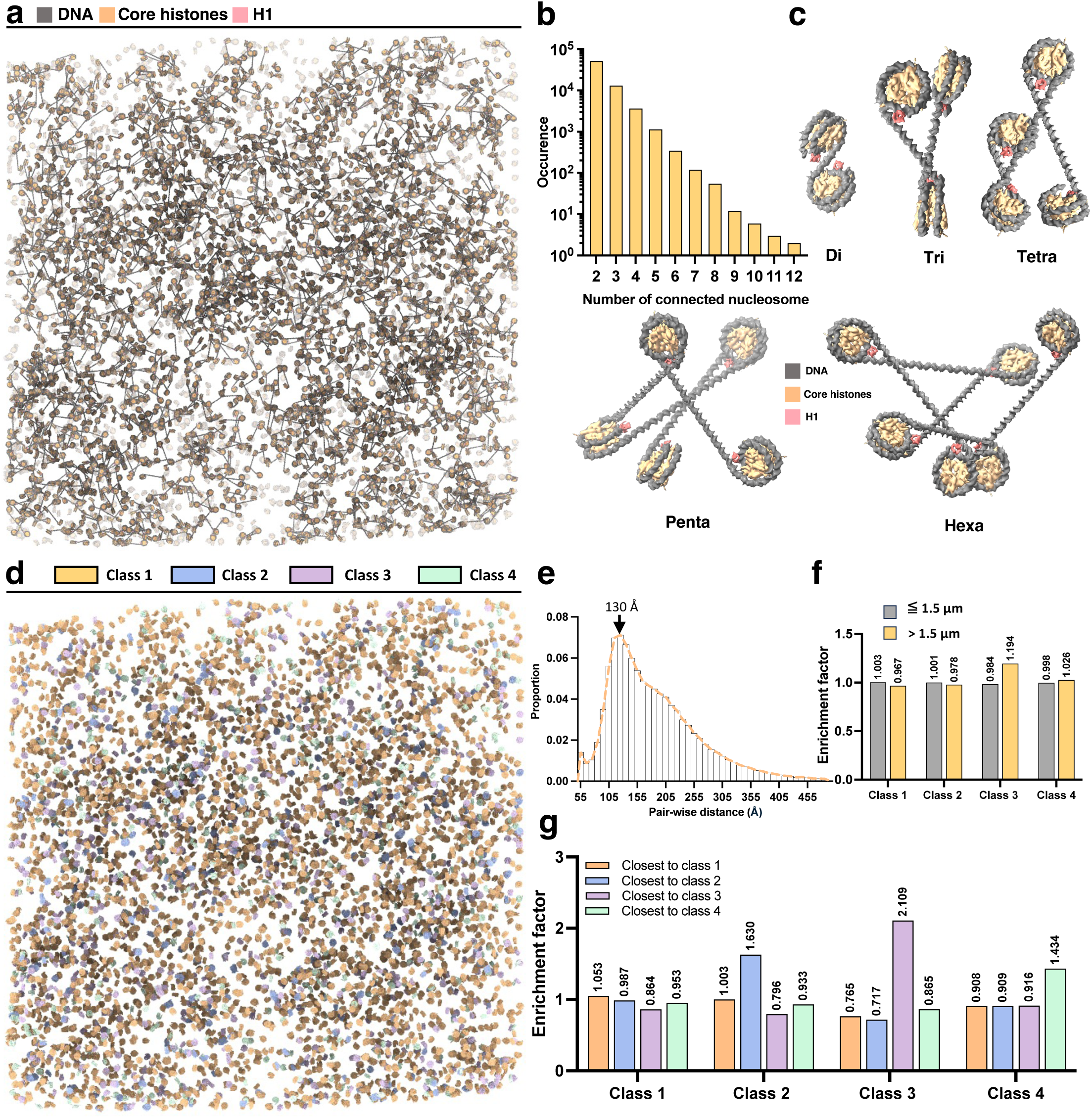
*In situ* chromatin architecture in the nucleus. **a**, A map-back representation of nucleosomes linked by linear DNA strand of various lengths in a tomogram collected from a heterochromatin region without HIV-1 cores. Only zig-zag linked nucleosomes (see methods section) are highlighted for clarity. **b**, A histogram showing the distribution of nucleosomes in various zig-zag connected oligomers (n = 253 tomograms). **c**, Representative zig-zag linked di-, tri-, tetra-, penta-, and hexa-nucleosomes. **d**, A map-back representation of four characterized nucleosome classes in a tomogram collected from a heterochromatin region without HIV-1 core. Nucleosome classes are colored accordingly. **e**, Pair-wise nucleosome distance distribution (n = 253 tomograms), the peak is indicated. **f**, The distribution of nucleosome classes in periphery (distance to NE ≦ 1.5 μm) (n = 235 tomograms) and interior (distance to NE > 1.5 μm) regions of nuclei (n = 18 tomograms). The enrichment factor is calculated by the radio of regional nucleosome proportion over the global proportion of each class. Two-sided Fisher’s exact test is applied for each class: for class 1, p = 1.8372 × 10^-11^ < 0.0001, n = 146,656 (≦ 1.5 μm), n = 11,780 (> 1.5 μm); for class 2, p = 0.3654 > 0.05, no significance, n = 17,168 (≦ 1.5 μm), n = 1,395 (> 1.5 μm); for class 3, p < 1 × 10^-15^, n = 22,844 (≦ 1.5 μm), n = 2,309 (> 1.5 μm); for class 4, p = 0.1721 > 0.05, no significance, n = 25,748 (≦ 1.5 μm), n = 2,206 (> 1.5 μm). **g**, The distribution of nearest neighbour classes (coloured) for each nucleosome class. The enrichment factor is calculated by the ratio of local (the nearest neighbor) nucleosome proportion over the global proportion of each class. Two-sided Chi-square test is applied for each class: for class 1, p < 1 × 10^-15^, n = 158,436; for class 2, p < 1 × 10^-15^, n = 18,563; for class 3, p < 1 × 10^-15^, n = 25,153; for class 4, p < 1 × 10^-15^, n = 27,954.

Consistent with our previous observations^48^, chromatin was not organized into regular 30-nm fibers, but instead exhibited a heterogeneous architecture, in which compact chromatin regions coexisted with relaxed, irregular zigzag arrangements of nucleosomes (Fig. 3a, Supplementary Fig. 6, 7a, Supplementary Video 4). We also observed chromatin loops and locally zig-zagged nucleosome clutches connected by visible linker DNA segments (Supplementary Fig. 6, 7a). To infer multi-nucleosome connectivity and linker geometry directly from tomograms, we developed a physics-based probabilistic linker-assignment framework, inspired by Kreysing et al^49^. Nucleosome coordinates and orientations derived from subtomogram refinement defined two DNA exit arms per particle. Candidate inter-nucleosome linkages were iteratively evaluated for geometric compatibility and assigned using a probability-ranked greedy algorithm, ensuring that each DNA arm participated in only one connection and that no closed cycles formed. The resulting connectivity graph revealed irregularly packed chromatin trajectories spanning kilobase scales, from local, short-range zigzag configurations to the longest continuous chains comprising up to 12 nucleosomes (∼2 kb of DNA) (Fig. 3b–c).

To examine whether specific nucleosome classes exhibit spatial preferences within the nucleus, we mapped all four classes of nucleosomes back into their original tomograms (Fig. 3d, Supplementary Fig. 7b) and quantified their global and local distributions relative to the nuclear envelope (≤1.5 µm vs. >1.5 µm from the periphery) (Fig. 3e, f). Pairwise nearest-neighbor distance analysis revealed a dominant inter-nucleosome spacing of ∼120–140 Å, accompanied by a secondary shorter-distance peak at ∼55 Å (Fig. 3e). These distances correspond to separations between i and i + 1 nucleosomes within zigzag chromatin fibers and between i and i + 2 nucleosomes in stacked H1-bound configurations, respectively. H1-bound nucleosomes (Class 1) were significantly enriched near the nuclear periphery (p = 1.8 × 10⁻¹¹), whereas core nucleosomes (Class 3) were preferentially localized to the nuclear interior (1.19-fold, p < 1 × 10⁻¹⁵). In contrast, both stacked H1-bound nucleosomes and open-linker H1-bound nucleosomes (Classes 2 and 4) exhibited relatively uniform distributions across peripheral and interior nuclear regions (Fig. 3f).

We next investigated whether different nucleosome classes exhibited preferential association patterns with themselves or with other classes. All four classes showed some degree of self-association, though to varying extents (Fig. 3g). Core nucleosomes (Class 3) displayed the strongest self-association with >2-fold enrichment, followed by stacked H1-bound nucleosomes (Class 2) with a 1.63-fold enrichment and open-linker H1-bound nucleosomes (Class 4) with a 1.43-fold enrichment. In contrast, H1-bound nucleosomes (Class 1) showed minimal preference among the four classes (Fig. 3g). These findings indicate that nucleosome organization within the nucleus is non-uniform yet non-random, suggesting the presence of domains enriched in specific chromatin states that collectively contribute to a dynamic and heterogeneous chromatin architecture. Further correlative cryo-ET analyses will be instrumental in elucidating which nuclear factors and histone/DNA modifications underlie the formation and regulation of these distinct chromatin domains, insights that are not readily captured by conventional bulk measurements.

### HIV-1 cores locally exclude nucleosomes and prefer open chromatin

To investigate the chromatin organization encountered by HIV-1 during nuclear traversal, we analyzed nucleosome distributions surrounding HIV-1 cores in tomograms. Nucleosome densities were projected through 130 nm thick lamellae to generate 2D heatmaps (Fig. 4a). These maps revealed that chromatin is less condensed in deeper nuclear regions compared to areas adjacent to the nuclear envelope, as expected. Consistent with previous reports^49, 50^, the regions directly beneath the NPC basket were consistently devoid of nucleosomes (Fig. 1e).

**Figure 4.**
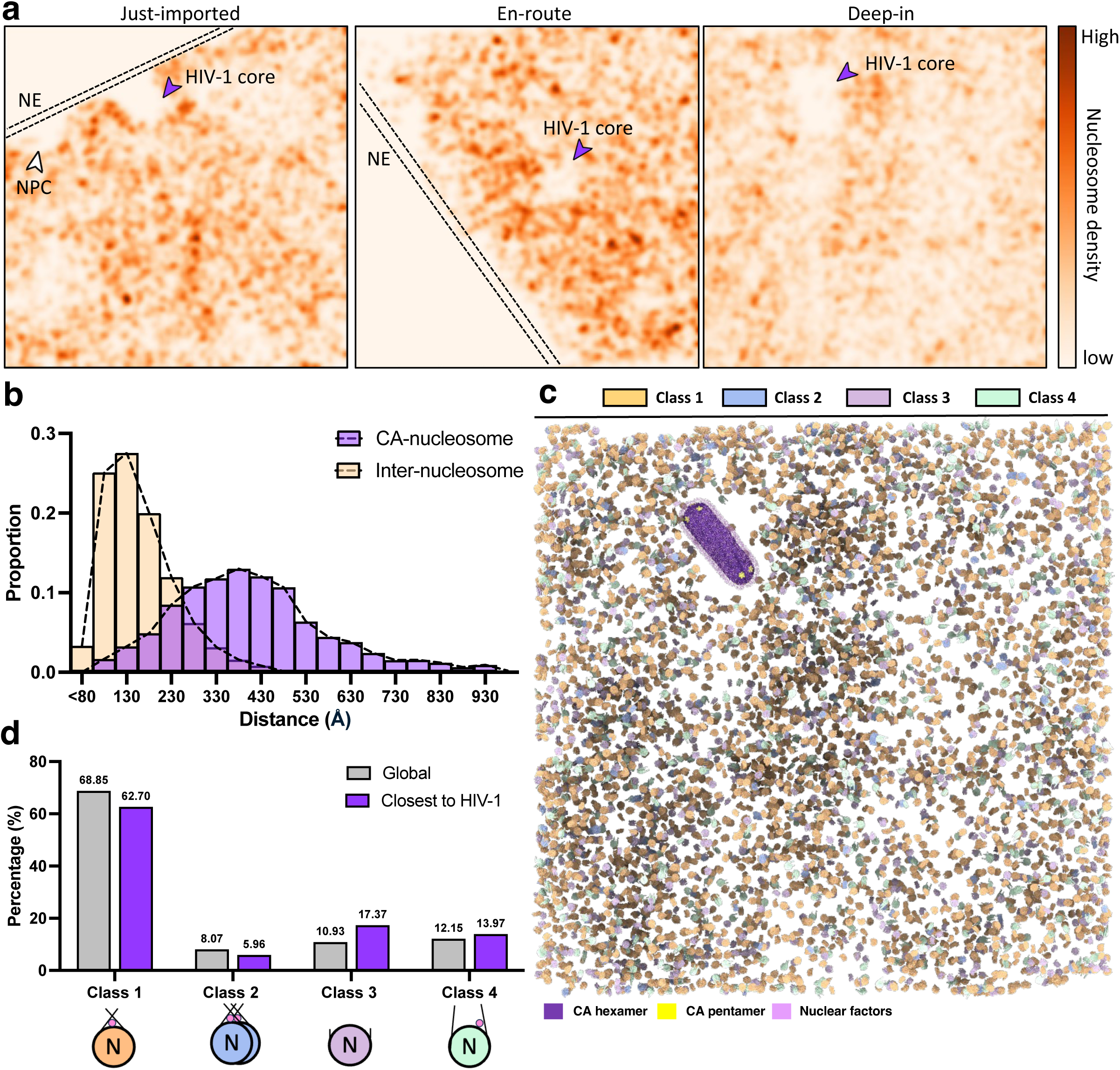
Chromatin exclusion by HIV-1 cores during nuclear trafficking. **a**, Heat maps from the three tomograms in Figure 1d-i, showing the map-backed nucleosome density (projected over consistent 130 nm thick tomogram) during HIV-1 nuclear traversal. Areas occupied by HIV-1 cores are indicated by purple arrowheads. NPC occupation is indicated by the white arrowhead. NE is indicated by the black dashed lines. **b**, The distributions of pair-wise inter-nucleosome distance and distance between HIV-1 CA and nucleosomes. **c**, Map-back of nucleosomes by classes in interior region of the nucleus containing a HIV-1 core, with all classes included and coloured accordingly. The HIV-1 core and nuclear factors are mapped back as in Figure 1i. **d**, The distribution of nucleosome classes in the nearest neighbours of HIV-1 cores compared to the global distribution. Two-sided Fisher’s exact test is applied: for class 1, p < 1 × 10^-15^, n = 2,787; for class 2, p = 1.14523030 × 10^-7^ < 0.0001, n = 265; for class 3, p < 1 × 10^-15^, n = 772; for class 4, p = 3.02791446817 × 10^-4^ < 0.001, n = 621.

Strikingly, we observed a persistent nucleosome-free zone in the immediate vicinity of HIV-1 cores, regardless of their position within the nucleus (Fig. 4a, n = 55 cores). Density profiling around the cores showed that while nuclear factors occupied regions approximately 7 nm from the capsid surface, the nucleosome-depleted zone extended up to ∼35 nm (Supplementary Fig. 9). Moreover, pairwise nearest-neighbor distance analysis revealed that typical inter-nucleosome distances were 120–140 Å, whereas capsid–nucleosome separations peaked around 350 Å (Fig. 4b). Together, these observations support a model in which HIV-1 cores locally exclude nucleosomes, potentially favoring access to open chromatin, although the mechanism of exclusion remains to be determined.

We next analyzed the spatial distribution of the four nucleosome classes surrounding HIV-1 cores by quantifying their local enrichment, defined as the ratio of each class’s proportion within the region of HIV-1 cores to its global nuclear proportion (Fig. 4c, d). This analysis revealed that HIV-1 cores strongly prefer open chromatin regions containing core nucleosomes (Class 3) and open-linker H1-bound nucleosomes (Class 4) (Fig. 4d). In contrast, stacked H1-bound nucleosomes (Class 2) were the least favored by HIV-1 cores (Fig. 4d). These findings suggest that HIV-1 cores selectively associate with accessible chromatin environments, potentially facilitating their trafficking toward nuclear speckle–associated domains (SPADs), where viral integration preferentially occurs.

### CPSF6 facilitates nuclear trafficking of HIV-1 cores

CPSF6 is a key nuclear host factor that dictates integration site selection^28, 51, 52^. It is thought to direct HIV-1 cores away from heterochromatin-rich regions toward the nuclear interior, thereby promoting integration into gene-dense, transcriptionally active chromatin within SPADs^29, 31, 53^. CPSF6 contains a single phenylalanine–glycine (FG) motif, and mutation of this motif abolishes its association with the HIV-1 capsid^53^, resulting in a loss of CPSF6-dependent targeting of integration into euchromatin and transcriptionally active genes, as demonstrated by integration-site mapping and fluorescence microscopy studies^27, 28^. Conversely, HIV-1 capsids carrying the N74D mutation fail to bind CPSF6^53^ and are defective in CPSF6-dependent trafficking to nuclear speckles and integration into SPADs^31, 54^.

We analyzed the nuclear trafficking of HIV-1 cores carrying the N74D mutation and compared them with WT cores. Although N74D cores exhibited similar morphologies and nuclear envelope binding as WT cores, they failed to accumulate efficiently within nuclei (Supplementary Fig. 1b, c). Instead, mNeonGreen-IN puncta were predominantly localized near the nuclear periphery and were rarely observed in the nuclear interior. Correlative cryo-ET imaging further revealed that most N74D nuclear cores remained tethered to NPCs, with only a single detached core detected near the nuclear envelope and none penetrating deeper into the nucleus (Fig. 5a-c, Supplementary Fig. 10, Supplementary Video 5). Notably, these N74D cores lacked the prominent surface densities observed coating imported WT cores (Fig. 5a-b). Density profiling confirmed the presence of distinct density layer at a distance of ∼7 nm from the capsid layer only around imported WT cores, but absent in both N74D cores within the nucleus and WT cores outside the nucleus (Fig. 5d), consistent with our previous observations^19^.

**Figure 5.**
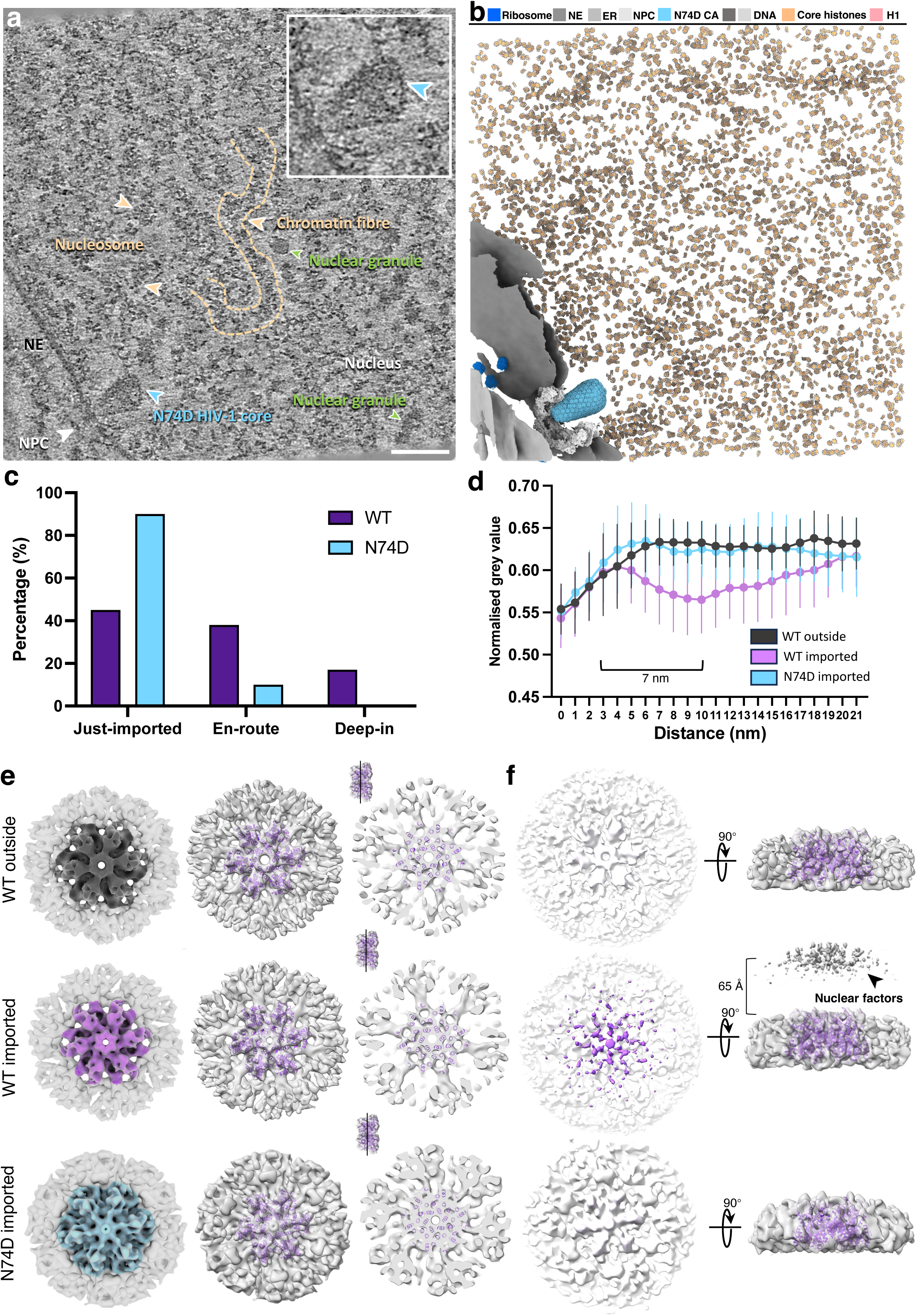
CPSF6 facilitates the nuclear traversal of HIV-1. **a-b,** A representative tomographic slice a “Just-imported” cone-shaped N74D core **(a)** and corresponding segmented volume **(b)**. No discernable core-surrounding density is observed (enlarged in the inset). The NPC, ribosomes, nucleosomes, nuclear granules, and chromatin fibres are labelled. The nucleus, NE, and ER are annotated accordingly. Ribosomes, NE, ER, N74D CA hexamers, DNA, core histones and histone H1 of nucleosomes are segmented and mapped back with the indicated colours. Of note, three subunits of the NPC exceeds the upper bound of tomogram, thus are not depicted. Scale bar, 100 nm. **c**, Distributions of WT and N74D cores in the nucleus. Fisher’s exact test is applied, p = 0.0424 < 0.05, n of WT cores = 55, n of N74D cores = 10. **d**, Density profiling of the vicinity of cores. The same strategy illustrated in **Supplementary Figure 9** is applied with shorter lines. For WT outside cores, n of lines = 155, n of cores = 12. For WT imported cores, n of lines = 111, n of cores = 10. For N74D imported cores, n of lines = 101, n of cores = 10. The central lines represent the mean, and whisks represent the S.D. The distance from the surface of imported WT HIV-1 cores to the density center of nuclear factors approximates 7 nm. **e**, Cryo-ET STA structures of the HIV-1 CA hexamers in capsid lattices from WT cores outside the nucleus (top), WT cores inside the nucleus (middle), and N74D cores inside the nucleus (bottom). The left panel: top views of the STA map (contoured at 2.5σ) with the central hexamer coloured; the middle panel: top views of the STA map (WT contoured at 2.5σ, N74D contoured at 1.5σ) fitted with a CA hexamer model (PDB 6SKK); the right panel: A slice of CA hexamer density maps (position shown in the upper left corner) fitted with atomic model (PDB 6SKK). **f**, CPSF6 density shown in the imported WT CA STA map (middle panel with an arrowhead), are absent in the STA maps of imported N74D cores (bottom) and outside WT cores (top). The left panel: CA hexamer density maps contoured at 0.25σ, coloured according to the height, from white (bottom) to purple (top); the right panel: the side views of CA hexamer density maps (major hexamer density contoured at 2.5σ for WT and 1.5σ for N74D, density of nuclear factors contoured at 0.25σ), the distance between CA hexamer and center of nuclear factors measures approximately 65 Å for the imported WT core.

To further characterize capsid-nuclear factor interactions, we performed STA of CA lattices from both WT and imported N74D cores which had not been previously resolved^19^. This analysis yielded three CA hexamer maps from WT cores outside the nucleus, WT cores inside the nucleus, and N74D cores inside the nucleus, resolved at 10.1 Å, 9.6 Å, and 12.8 Å, respectively (Fig. 5e, Supplementary Fig. 11, Supplementary Table 3). Only imported WT cores, but not unimported WT or N74D cores, exhibited additional densities ∼65 Å above the capsid surface (Fig. 5e-f). The **location** of these densities is consistent with structural studies showing the prion-like low-complexity regions (LCRs) of CPSF6 bound to CA tubes^55^, providing direct *in situ* evidence that CPSF6 interacts with the viral capsid, likely guiding HIV-1 core trafficking from peripheral heterochromatin toward the transcriptionally active nuclear interior.

## Discussion

For decades, determining the *in situ* structures of chromatin and nucleosomes within intact nuclei has remained a major challenge. Recent advances in cryo-FIB milling and cryo-ET have made it possible to visualize native chromatin fibers and nucleosomes at moderate resolutions (> 10 Å)^48, 56, 57, 58, 59^. In this study, we resolved the native nucleosome structure at 5.6 Å resolution, providing an unprecedented level of structural detail for a ∼250 kDa complex within the crowded environment of the cell nucleus. Our analyses further revealed multiple nucleosome classes corresponding to compact and open chromatin conformations, and distinct histone tail organizations critical for epigenetic regulation and other DNA metabolic processes. *In situ* visualization of off-dyad H1 binding uncovered asymmetric linker DNA engagement, providing structural insight into the flexibility and dynamic behavior of chromatin. These findings mark a step change in the direct visualization of chromatin architecture and heterogeneity within intact nuclei.

HIV-1 post-nuclear events have been extensively investigated through cell biology approaches^28, 32, 51, 60^, fluorescence imaging^61, 62, 63, 64, 65^, and more recently, cryo-ET^19, 23, 24, 25^. These studies established that HIV-1 enters the nucleus with a largely intact capsid, which interacts with CPSF6 on the nuclear side and is subsequently directed toward SPADs, where integration preferentially occurs within transcriptionally active chromatin. However, how viral cores navigate the crowded nuclear environment and complex chromatin landscape to reach these integration sites has remained unclear. By combining a functional nuclear import system with high-resolution *in situ* structural analysis, we directly imaged HIV-1 cores at varying depths of nuclear penetration and characterized their interactions with chromatin inside intact nuclei.

A key finding is that HIV-1 cores exhibit distinct preferences for specific local chromatin environments. Viral cores excluded nucleosomes from within ∼35 nm of their capsid surface, creating local nucleosome-depleted zones. Similar exclusion zones were observed beneath NPCs, both in this study and by others^49, 50^, and are thought to facilitate macromolecular transport across the nuclear envelope. Within the surrounding chromatin, open nucleosome states (Classes 3 and 4) were enriched, whereas compact nucleosome states (Classes 1 and 2) were underrepresented. This pattern suggests that nuclear factor–decorated HIV-1 cores preferentially traverse open, accessible chromatin regions, potentially displacing nucleosomes as they progress toward transcriptionally active integration sites.

We further show that CPSF6 decorates the HIV-1 core upon nuclear entry, enabling the viral core to traverse compact peripheral chromatin and access the more dispersed chromatin of the nuclear interior. Disruption of the CA–CPSF6 interaction (as in the N74D mutant) confined viral cores to the nuclear periphery, consistent with previous observations that loss of CPSF6 association redirects HIV-1 integration toward lamina-associated heterochromatin^28, 30, 31, 65, 66, 67, 68^. The C-terminal arginine/serine-rich domain of CPSF6 is intrinsically disordered and capable of mediating phase separation and nuclear speckles association^60, 69^. Whether this condensation property contributes to the nuclear trafficking of HIV-1 cores toward nuclear speckle remains to be determined. Future studies should also explore whether additional nuclear factors cooperate with CPSF6 to regulate HIV-1 trafficking and integration site selection.

Based on these findings, we propose a model in which CPSF6 binding and chromatin accessibility act in concert to guide HIV-1 cores toward transcriptionally active regions (Fig. 6). Upon nuclear entry, HIV-1 cores rapidly recruit CPSF6 and potentially other nuclear cofactors, forming a proteinaceous coat on the viral capsid that facilitates passage through compact peripheral chromatin into the more open and accessible chromatin of the nuclear interior. During this process, HIV-1 cores locally exclude nucleosomes through a yet-unknown mechanism.

**Figure 6.**
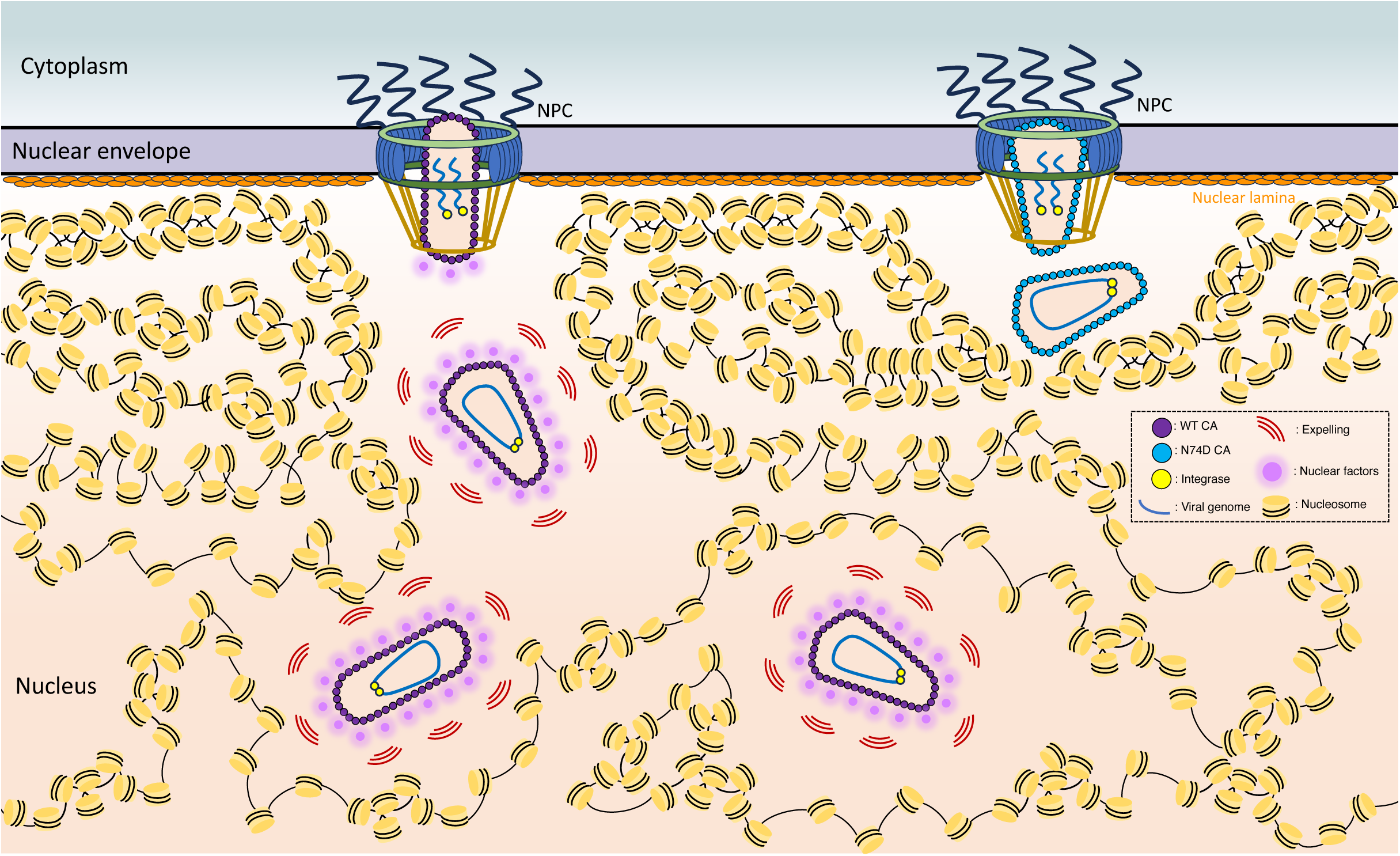
Schematic model of HIV-1 nuclear trafficking. This schematic summarizes the distinct behaviors of WT and CPSF6-binding-deficient (N74D) HIV-1 cores during the nuclear trafficking. Upon nuclear import, WT HIV-1 cores rapidly engage nuclear factors such as CPSF6, which assemble into a phase-separation-like layer that excludes chromatin along the trajectory of core traversal. This CPSF6-associated environment facilitates HIV-1 core movement through the condensed heterochromatin at the nuclear periphery. As the cores traverse into the nuclear interior, they encounter regions of less-compact chromatin enriched in dynamically positioned nucleosomes, likely corresponding to transcriptionally active domains. In contrast, N74D HIV-1 cores, which fail to interact with CPSF6 and related nuclear factors, are unable to penetrate the dense peripheral heterochromatin and consequently remain trapped at the nuclear periphery.

Collectively, these findings establish a structural framework directly linking HIV-1 nuclear trafficking to chromatin organization, providing a foundation for future studies of integration site selection and viral latency. More broadly, this work highlights the potential of *in situ* structural biology to illuminate the 3D organization of the genome, as well as the mechanisms of replication, repair, and transcriptional regulation across physiological and disease contexts.

## Methods

### Mammalian cell culture

All cell cultures were maintained at 37 °C in a humidified incubator with 5% CO₂. Human embryonic kidney (HEK) 293T Lenti-X cells (632180; Takara/Clontech) were obtained directly from the vendor, who performed authentication by short tandem repeat (STR) profiling and confirmed the absence of mycoplasma contamination. HEK293T Lenti-X cells were grown in Dulbecco’s modified Eagle medium (DMEM; Gibco) supplemented with 10% fetal bovine serum (FBS), 2 mM L-glutamine (Gibco), and 1% MEM non-essential amino acids (Gibco). CD4⁺ T lymphocyte CEM cells (NIH HIV Reagent Program, ARP-117) were maintained in RPMI-1640 medium (Sigma-Aldrich) containing 10% FBS and 2 mM L-glutamine (Gibco). These cells were not independently authenticated following acquisition.

### Production of mNeonGreen-IN-labelled HIV-1 cores containing near-full-length HIV-1 RNA

mNeonGreen-IN–labelled HIV-1 cores carrying near–full-length viral RNA were generated as previously described^19^. Briefly, HEK293T Lenti-X cells (∼80% confluence, four T175 flasks) were transfected using polyethyleneimine (PEI; branched, MW ∼25,000; Sigma-Aldrich). Each transfection contained 24 µg of Env-defective HIV-1 plasmid (R9ΔE-CA-WT or R9ΔE-CA-N74D) and 6 µg of Vpr–mNeonGreen–IN plasmid (a gift from Prof. Zandrea Ambrose, University of Pittsburgh) mixed with 120 µg PEI in Opti-MEM. After incubation at room temperature for 20 min, the mixture was added to the cells. 16 h post-transfection, the medium was replaced with fresh DMEM supplemented with 10% FBS, 1% nonessential amino acids, and 1% L-glutamine. Supernatants containing HIV-1 particles were collected after 24 h.

Virus-containing supernatants (120 ml) were filtered through a 0.45 µm filter (Sarstedt) and layered onto 5 ml of 20% sucrose in 1× STE buffer (10 mM Tris-HCl pH 7.4, 100 mM NaCl, 1 mM EDTA) in four 38.5 ml ultracentrifuge tubes (Beckman Coulter). Virions were pelleted using a SW32Ti rotor at 85,500 × g for 3 h at 4 °C (Optima XPN-90, Beckman Coulter). Each pellet was resuspended in 400 µl 1× STE supplemented with 1 mM IP6.

For core purification, a continuous sucrose gradient (30–85%) was prepared in 13.2 ml ultracentrifuge tubes (Beckman Coulter) in 1× STE with 1 mM IP6. The gradient was overlaid with 250 µl of 15% sucrose containing 1% Triton X-100, followed by 7.5% sucrose to create a protective barrier. Approximately 800 µl of concentrated HIV-1 particles was layered on top, and gradients were centrifuged at 182,600 × g for 16 h at 4 °C using a SW41Ti rotor.

After centrifugation, the fluorescent mNeonGreen band corresponding to HIV-1 cores was visualized under blue light and collected by side puncture with a 25 G needle (BD Microlance 3). Core fractions were quantified by p24 ELISA and either snap-frozen in single-use aliquots at −80 °C or dialyzed overnight at 4 °C against 500 ml of 1× SHE buffer (10 mM HEPES-NaOH pH 7.4, 100 mM NaCl, 1 mM EDTA) containing 0.8 mM IP6 using a 7 kDa cut-off Slide-A-Lyzer Dialysis Cassette (Thermo Fisher Scientific) before mixing with permeabilized T cells.

### T cell membrane permeabilization

Exponentially growing CEM T cells (∼1 × 10⁶ cells/ml) were collected by centrifugation at 500 × g for 5 min at 4 °C, washed twice with 10 ml of ice-cold PBS, and pelleted under the same conditions. The cell pellet was resuspended in lysis buffer (20 mM HEPES pH 7.5, 25 mM KCl, 5 mM MgCl₂, 1 mM DTT, and 1× protease inhibitor) containing 0.018% digitonin. Cells were gently rotated at room temperature for 10 min to permeabilize the plasma membrane, following previously established procedures^19^. After incubation, the cells were briefly centrifuged at 200 × g for 1 min, and the supernatant was discarded. The pellet was resuspended in fresh lysis buffer without digitonin to remove residual detergent. To verify nuclear integrity, digitonin-treated CEM cells were incubated with 0.2 mg/ml FITC–dextran (500 kDa; Sigma-Aldrich, 46947) in lysis buffer for 15 min at room temperature. Images were acquired using a Leica TCS SP8 confocal microscope.

### Nuclear import of mNeonGreen-IN-labelled HIV-1 cores in permeabilized T cells

Nuclear import of mNeonGreen-IN-labelled HIV-1 cores in permeabilized T cells was carried out as previously described^19^. Nuclei of digitonin-permeabilized CEM cells were stained on ice with 1 µM SiR-DNA (Spirochrome) for 30 min. Approximately 4 × 10⁵ permeabilized cells were then incubated with HIV-1 cores (WT or N74D) at a final CA concentration of 20 µg/ ml in a 40 µl reaction mixture containing 0.2 mM IP6, 20 mM HEPES (pH 7.5), 18.8 mM KCl, 20 mM NaCl, 0.25 mM EDTA, 3.8 mM MgCl₂, 1 mM DTT, and 1× protease inhibitor cocktail. After a 30 min incubation on ice, 10 µl of supplement mix (rabbit reticulocyte lysate + ATP + GTP + creatine phosphate + creatine kinase) was added, resulting in final concentrations of 16% (v/v) rabbit reticulocyte lysate (RRL; Promega, L4151), 0.6 mM ATP, 0.06 mM GTP, 5 mM creatine phosphate, and 10 U µl⁻¹ creatine kinase. The reaction was incubated at 37 °C for 30 min to allow nuclear import. For confocal microscopy, samples were transferred to µ-Slide 18-well glass-bottom chambers (Ibidi) and imaged using a Leica TCS SP8 confocal microscope. For cryo-FIB milling and cryo-ET, samples were fixed with 5 mM ethylene glycol bis(succinimidyl succinate) (EGS) for 30 min at room temperature, quenched with 50 mM Tris-HCl (pH 7.5), and kept on ice until grid preparation.

### Confocal microscopy and fluorescence image processing

All fluorescence imaging was performed using a Leica TCS SP8 confocal microscope equipped with a HC PL Apo 63×/1.2 NA MotCORR water-immersion CS2 objective and a HyD GaAsP detector, operated through LAS X software (Leica Microsystems). Fluorescent signals from GFP and SiR-DNA were acquired using 488 nm and 633 nm excitation laser lines, respectively, with optimized spectral filters to minimize channel overlap. Z-stacks were acquired with a step size of 0.3–0.5 µm using a 1 a.u. pinhole and an image resolution of 1,024 × 1,024 pixels. Image stacks were processed and analysed using Leica Application Suite X (LAS X) and Fiji (ImageJ)^72^.

### Plunge-freezing vitrification

Sample vitrification was performed using a Leica EM GP2 automated plunge freezer under controlled conditions (20 °C, 95% relative humidity). The mixture of HIV-1 cores and permeabilized CEM cells was incubated with 10% glycerol for 2 min prior to freezing. A 3–6 µL aliquot of the mixture (containing approximately 4,000 cells per µL) was applied to the carbon side of glow-discharged Quantifoil R2/1 Cu 300 grids, while 1 µL of PBS was placed on the reverse side. Grids were back-blotted for 6 s using Whatman filter paper and then rapidly plunge-frozen in liquid ethane. For HIV-1 cores, 3.5 µL of WT or N74D cores were applied to the carbon side of glow-discharged lacey carbon film grids (Cu 300 mesh, Electron Microscopy Sciences), back-blotted for 3 s, and plunge-frozen in liquid ethane under identical environmental conditions. Cryo-EM data were collected on a Glacios transmission electron microscope (Thermo Fisher Scientific) equipped with a Falcon III direct electron detector at a nominal magnification of 73,000×.

### Correlative cryo-FIB milling

Vitrified mixtures of permeabilized CEM cells supplemented with RRL-ATP and HIV-1 cores were thinned by cryo-FIB milling to produce lamellae, guided by cryo-CLEM using two distinct systems. Two grids of WT cores were loaded via an Autoloader onto a dual-beam FIB/SEM microscope (Arctis, Thermo Fisher Scientific), equipped with a cryogenic stage cooled to −191 °C, a wide-field integrated fluorescence microscope (iFLM; 100×, NA 0.75), and a plasma multi-ion source (argon, xenon, and oxygen). Argon was used as the FIB source in this study.

Prior to milling, a platinum layer was sputter-coated for 30 s, followed by deposition of an organometallic platinum layer via the GIS system (Thermo Fisher Scientific) for 50 s. Three-dimensional correlative milling was performed using the embedded protocol in WebUI v1.3 (Thermo Fisher Scientific) with a milling angle of 10°. For each position, a 15 µm z-stack of fluorescence (GFP) and reflection images was acquired at 500 nm step size after rough milling under default parameters. Target fluorescence coordinates were registered using discernible ice chunks as fiducial markers in both SEM and FIB images to guide lamella placement. Lamellae were produced stepwise: (i) opening at 2 nA, (ii) rough milling at 0.74 nA and 0.2 nA, and (iii) polishing at 60 pA, yielding a final lamella thickness of 140 nm. To enhance fluorescence detection on polished lamellae, the step size was reduced to 100 nm, generating a 15 µm stack (101 images) in both GFP and far-red channels.

Four grids of WT cores and one grid of N74D cores were milled on a dual-beam FIB/SEM microscope (Aquilos 2, Thermo Fisher Scientific) equipped with a cryo-transfer system and a rotatable cryo-stage cooled to −191 °C. The instrument was modified to incorporate a fluorescence module (METEOR, Delmic Cryo BV; 50×, NA 0.8). Grids were mounted on a METEOR shuttle with a pre-tilt of 26° and coated with an organometallic platinum layer for 30 s using the GIS system. Milling was performed at 10° using AutoTEM 5 (Thermo Fisher Scientific). Cells located near grid centers were selected for lamella preparation. Before milling, fluorescence stacks were acquired for each site in GFP and far-red channels at 200 nm steps (± 6 µm), processed in ImageJ to improve signal-to-noise ratio (SNR), and correlated with FIB/SEM images using the 3D Correlation Toolbox^73^, with ice features as fiducial markers. Sequential milling was performed from 0.5 nA (rough) to 0.3 nA (medium), 0.1 nA (fine), 60 pA (first polishing), and 30 pA (final polishing), yielding lamellae ∼120 nm thick. Final fluorescence stacks were collected (GFP and far-red) at 100 nm steps (± 2 µm) using a 400 mW laser and 300 ms exposure. In total, 15 lamellae (Arctis) and 21 lamellae (Aquilos 2) containing detectable fluorescence were generated for WT cores, and 9 lamellae were produced in Aquilos 2 for N74D cores.

### Cryo-electron tomography data collection

Lamellae were transferred to two Titan Krios G3 (Thermo Fisher Scientific) microscopes operating at 300 kV, each equipped with a Falcon 4i detector and Selectris X energy filter. Objective apertures (100 µm) were inserted. Overview images were collected at 8100× magnification and stitched to create lamella maps, which were aligned with corresponding fluorescence images in ImageJ. Tilt series were then collected at the correlated sites and neighboring regions (without overlap) using Tomography 5 (Thermo Fisher Scientific) at 64k× magnification.

For WT cores, 253 tilt series were recorded at 1.94 Å/pixel, using lamella pre-tilts of ± 12°, and a dose-symmetric scheme from −42° to +66° in 2° increments. Each series comprised 55 tilts with 5 movie frames per tilt at 2.5 e⁻/Å²/tilt (total dose 137.5 e⁻/Å²) and −2 to −5 µm defocus. For N74D cores, 45 tilt series were collected at 1.903 Å/pixel using identical imaging parameters.

### Alignment of tilt series

Movie frames were aligned with MotionCor2^74^. Tilt-series alignment and tomogram reconstruction were performed in IMOD v4.11^75^ by patch tracking (200×200 pixel patches, 0.45 overlap). Alignments were inspected manually and poor frames removed. Reconstructed tomograms were binned 6× (11.64 Å/pixel for WT, 11.418 Å/pixel for N74D) and SIRT-like filtered (8 iterations) for visualization.

### Density profiling

To measure the intensity of surround densities of HIV-1 cores, grey value measurements were taken to assess the change in density as a function of distance from the surface of HIV-1 cores. For each core, lines were derived from the capsid surface including partial density of the CA (∼ 3 nm). Black density was assigned a value of 0, and white was assigned a value of 1, with higher densities corresponding to lower values. In total, 233 lines were derived from 12 imported WT cores, 155 lines were derived from 12 outside WT cores, and 101 lines were derived from 10 imported N74D cores. For chromatin, 227 lines were derived and profiled in 12 tomograms.

### Template matching

To localize nucleosomes, template matching was performed using emClarity version 1.5.0.2^76^ with non-CTF-corrected tomograms binned at 6×. The procedure utilized a template derived from EMD-3947^77^, which was low-pass filtered to 40 Å (Supplementary Fig. 3a). An exhaustive search was applied by giving a peak threshold ranging from 1,500 to 3,500 based on the size of nuclear region in the tomogram. Matched coordinates were then imported into Chimera using the plug-in Place Object^78, 79^ for the initial cleaning based on the cross-correlation coefficient (ccc) value, the cut-off was set to 0.35. Particles residing outside nuclei or with a ccc value lower than 0.35 were treated as false positives and removed (Supplementary Fig. 3a).

To localize individual CA hexamers, template matching was performed in emClarity version 1.5.0.2^76^ with non-CTF-corrected tomograms binned at 6× using a low-pass filtered (30 Å) template derived from EMD-12452^80^. An exhaustive search was applied by giving a peak threshold of 2,000 in the small cropped-out tomogram region (150 nm^3^) containing the HIV-1 core. HIV-1 CA hexamer peaks were selected with the MagpiEM tool (available at https://github.com/fnight128/MagpiEM), and particle selection was identified based on the geometric constraints of the capsid lattice. Any hexamers that did not conform to the expected hexagonal lattice geometry were automatically excluded, followed by manual inspection to ensure the selection precision (Supplementary Fig. 11a).

NPCs and 80S ribosomes were located adopting the same procedure, using 1/8 of the original density map EMD-11967^23^ for NPCs, and EMD-1780^81^ for ribosomes, respectively.

### Subtomogram averaging

The 3D alignment and averaging of nucleosomes were conducted in RELION version 4.0^82^ with an initial particle number of 551,565. Prior to the iterative refinement, iterative classification was performed to further clean the particles until no prominent bad class was generated, resulting in 230,106 particles for the final classification (Supplementary Fig. 3a). In the final classification, four major classes were generated: class 1 (H1-bound nucleosome) with 158,436 particles, class 2 (stacked H1-bound nucleosomes) with 18,563 particles, class 3 (core nucleosome) with 25,153 particles, and class 4 (open-linker H1-bound nucleosome) with 27,954 particles. Iterative refinement was then carried out for each class from bin6 to bin1. The resolution of the reconstructed structure was calculated using a gold-standard Fourier shell correlation (FSC) cut-off of 0.143. For nucleosomes, the final resolution was determined at 5.6 Å, 20.0 Å, 9.9 Å, 9.3 Å, and 12.1 Å for class 1, class 2, mono H1-bound nucleosome in class 2, class 3, and class 4, respectively (Supplementary Fig. 3b). For HIV-1 CA hexamers, a similar procedure was employed (Supplementary Fig. 11a), and the resolution was determined at 10.1 Å, 9.6 Å, and 12.8 Å for outside WT CA (6,404 particles), imported WT CA (4,445 particles), and imported N74D CA (988 particles) (Supplementary Fig. 11b). Structural fitting was conducted in ChimeraX^83^.

### Segmentation

To enhance the segmentation, reconstructed bin6 tomograms were corrected for the missing wedge and denoised using IsoNet version 0.2^84^, applying 35 iterations with sequential noise cut-off levels of 0.05, 0.1, 0.15, 0.2, and 0.25 at iterations 10, 15, 20, 25, and 30, respectively. Membranes and nuclear envelopes in all tomograms were initially segmented using MemBrain-seg^85^, then imported into ChimeraX for manual cleaning and polishing. Nucleosomes mapped back to the tomograms with segmented membranes using ChimeraX and ArtiaX^86^, based on their positions and orientations after final refinement. For better visualization, nucleosomes depicted in the segmented volume were low-pass filtered structures from this study. HIV-1 cores were mapped back based on the refined coordinates and orientations of CA hexamers from subtomogram averaging, with missing CA hexamers manually placed back using the *in situ* CA hexamer structures resolved in this study. CA pentamers were only placed back on cores with most CA hexamers matched and the five surrounding hexamers identified and then modelled using the low-pass filtered EMD-13422^80^. NPCs and 80S ribosomes were mapped back based on their coordinates after the template matching using the low-pass filtered model generated from EMD-11967 and EMD-1780, respectively. Movies were made using ChimeraX.

### Definition of linker arms and geometric parameters

Each nucleosome was represented by two exit points (“linker arms”) and corresponding tangent vectors indicating the directions of DNA emerging from the dyad axis. The spatial positions and orientations of these linker arms were derived from subtomogram-averaged nucleosome densities and guided by the canonical chromatosome structure (PDB 7K5Y)^6^.

### Candidate pair identification

For each nucleosome, potential partners were identified based on the spatial proximity of linker-arm exit points. Neighbours located within 30 nm of either arm were considered candidate partners. For each valid pair (*i*, *j*), the four possible arm-to-arm combinations (arm₁–arm₁, arm₁–arm₂, arm₂–arm₁, arm₂–arm₂) were evaluated.

### DNA bending-energy model and connection probability

For each possible connection, the linker bending angle (θ) and arc length (L) were derived from the positions and orientations of the two arms, where θ is defined as the angle between the unit tangent vectors of the two arms, *t*_1_ and *t*_2_:

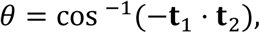

and L was computed from the distance between the two DNA exit points (ǁr₂−r₁ǁ) and the bending angle using circular-arc geometry:

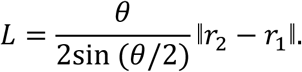

Where *r*_1_ and *r*_2_ are the coordinates of the two arms.

The DNA linker bending energy was then estimated using the worm-like chain (WLC) model, which describes the entropic elasticity of double-stranded DNA^87, 88, 89^:

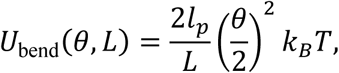

Where L is the linker length, *l*_*p*_= 50 nm, the persistence length of B-form DNA under physiological ionic conditions ^90^, and *k*_*B*_*T* is the thermal energy. The total connection probability was defined as:

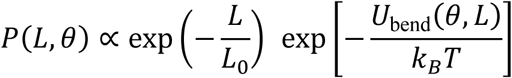

with *L*_0_ = 25 nm, the characteristic decay length. This formulation penalizes both excessively long and sharply bent linkers, thereby favouring geometrically plausible DNA trajectories.

### Probabilistic connection assignment and graph construction

A three-dimensional probability matrix *P*_*ijk*_ was computed, representing the likelihood of each possible arm-to-arm connection between nucleosome pairs (*i*, *j*), where *k* indexes the four possible arm combinations. Candidate connections were incorporated iteratively in descending order of probability.

At each iteration, the highest-probability entry was examined; if *P*_*ijk*_ > 0.1, the corresponding connection was accepted. A union–find (disjoint-set) data structure was used to track connected components, ensuring that each arm was used only once and that no cyclic connections occurred. After each accepted connection, all entries involving the used arms were excluded from further consideration. The process continued until no remaining probabilities exceeded the threshold. The resulting adjacency graph *G*(*V*, *E*) contains nucleosomes as vertices *V* and their most probable linker connections as edges *E*. Each connected component of *G* represents a continuous chromatin fragment of variable length and serves as the basis for further statistical and structural analyses.

### Statistical analysis

To assess the significance of distribution differences across nucleosomes, two-sided Chi-square test and Fisher’s exact test were applied. The calculation and plots were carried out using Prism v10.

## Data availability

All data needed to evaluate the conclusions in the paper are present in the paper and/or the Supplementary Information. The subtomogram-averaged maps for nucleosomes subunits have been deposited in the public database EMDB under the following accession codes: EMD-55446 (H1-bound nucleosome), EMD-55447 (stacked H1-bound nucleosomes), EMD-55448 (H1-bound nucleosome in stacking nucleosomes), EMD-55449 (core nucleosome), and EMD-55450 (open-linker H1-bound nucleosome). The subtomogram-averaged maps for the CA hexamers from both unimported and imported WT cores and from imported N74D cores have been deposited in the public database EMDB under the following accession codes: EMD-55441, EMD-55443, and EMD-55445, respectively. Source data are provided with this paper.

## Code availability

The scripts used in this study and relevant codes are deposited in GitHub (https://github.com/fnight128/MagpiEM).

## Acknowledgements

We thank L. Carrique, H. Duyvesteyn, D. Owen, J. Bancroft, E. Drydale and J. Gilchrist for their support in data collection. We thank J. Shi and C. Aiken (Vanderbilt University Medical Center) for kindly providing plasmids to produce HIV-1 cores. We thank F. Nightingale for the MagpiEM software. Y.S. is further supported by a CIHR fellowship from the Canadian Institutes of Health Research (194032), and an EMBO fellowship (ALTF 96-2024). We acknowledge the Oxford Particle Imaging Centre (OPIC) for access to cryo-FIB/SEM instruments (Arctis and Aquilos 2) and the cryo-EM instrument (Krios). We acknowledge the Oxford Cellular Imaging Core Facility (CICF) for access to fluorescence microscopes and imaging analysis software. We acknowledge Diamond Light Source for access to and support of the cryo-EM facilities at the UK national electron Bio-Imaging Centre (eBIC), proposal NT29812. Computation was performed at the Diamond Light Source and Oxford Biomedical Research Computing (BMRC) facility supported by the Wellcome Trust Core Award grant number 203141/Z/16/Z with additional support from the NIHR Oxford BRC. This work was supported by US National Institutes of Health grants U54AI170791 (P.Z.), R21AI184080 (P.Z.); the UK Wellcome Investigator Award 206422/Z/17/Z (P.Z.); the UK Wellcome Discovery Award 311427/Z/24/Z (P.Z.); the UK Biotechnology and Biological Sciences Research Council grant BB/S003339/1 (P.Z.); ERC AdG grant 101021133 (P.Z.); and the Chinese Academy of Medical Sciences (CAMS) Innovation Fund for Medical Science (CIFMS), China (grant no. 2024-I2M-2-001-1; P.Z., D.I.S.). The funders had no role in study design, data collection and analysis, decision to publish, or preparation of the paper.

## Contributions

P.Z. conceived the research. Z.H., Y.S. and P.Z. designed the experiments. Y.S. prepared samples. Y.S. conducted the fluorescence microscopy. J.S. helped with HIV-1 core preparation. Z.H. performed the cryo-CLEM and cryo-FIB. Z.H. and Y.S. collected the cryo-ET data and performed reconstruction of tomograms. Y.S. collected the micrographs of input cores. Z.H. performed STA and the segmentation. L.C. performed the linker DNA analysis under the supervision of D. I. S.. Z.H. performed the statistical analyses. Z.H. and Y.S. prepared figures. Z.H., Y.S., and P.Z. wrote the paper with help from other co-authors.

