## Supplementary figures and tables for "*In situ* chromatin dynamics and HIV-1 nuclear trafficking"

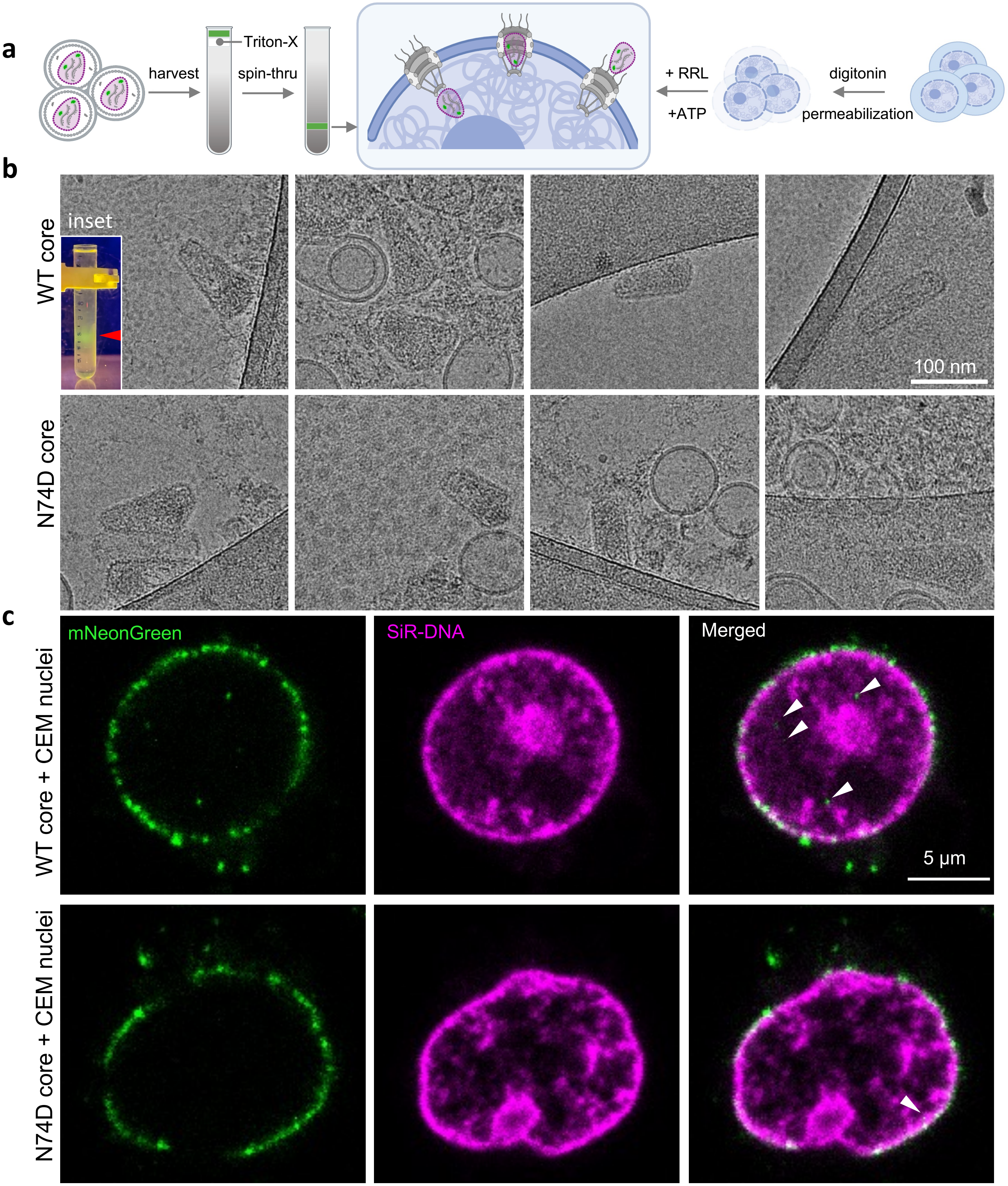


**Supplementary Figure 1 | Reconstitution of functional HIV-1 nuclear import using HIV-1 cores and permeabilized CEM cells.** **a**, Schematic illustration of HIV-1 cores and permeabilized T cell preparations. **b**, Representative electron micrographs of isolated HIV-1 cores from WT (top) and N74D variants (bottom). Scale bar, 100 nm. Inset, mNeonGreen-IN–labeled HIV-1 core band after 'spin-thru' detergent treatment, indicated by a red arrowhead. **c**, Representative confocal images of permeabilized CEM cells incubated with WT (top) or N74D (bottom) cores in the presence of RRL-ATP. Single z-slices from z-stacks are shown. HIV-1 cores are labeled with mNeonGreen-IN (green), and nuclei with SiR-DNA (magenta). Arrows indicate mNeonGreen-IN signals inside nuclei. Scale bar, 5 $\mu$m. Panel **a** was created with BioRender.com.

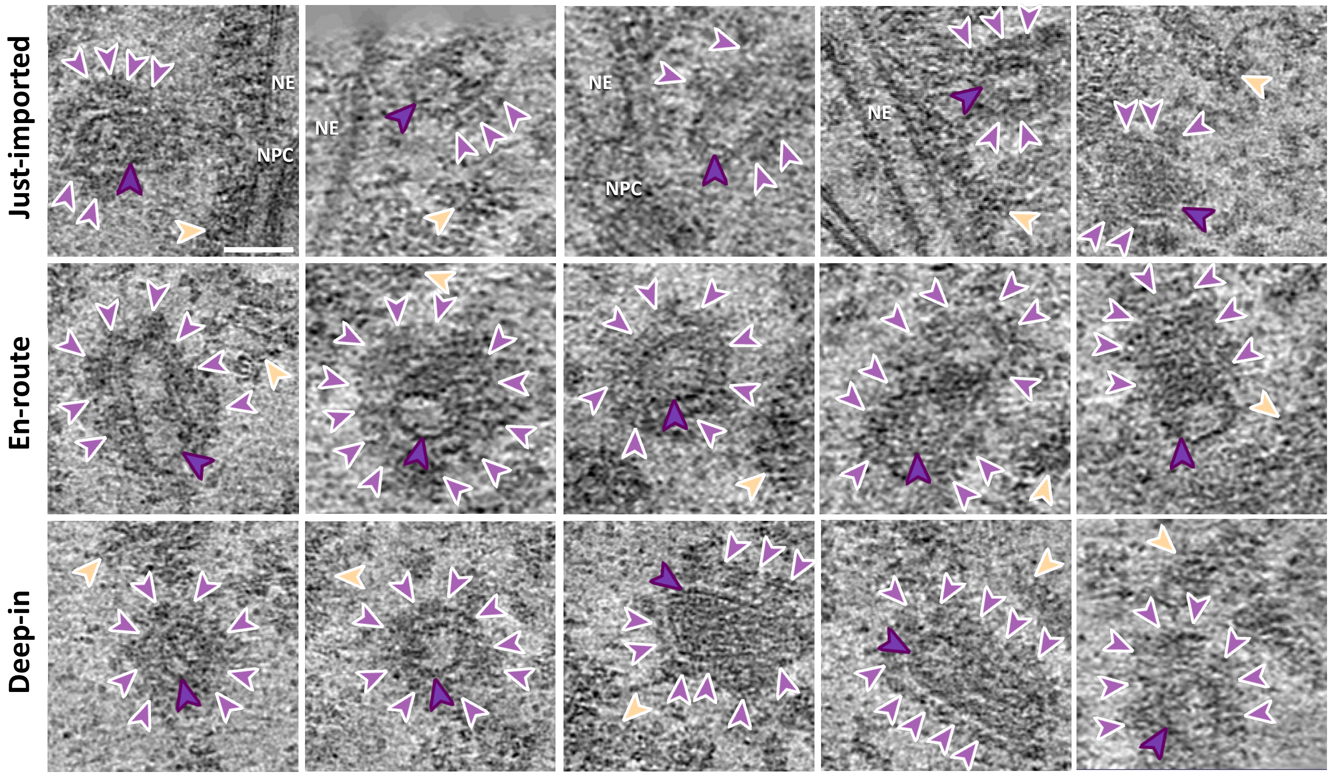


**Supplementary Figure 2 | Gallery of imported WT HIV-1 cores captured by correlative cryo-ET.** WT cores are indicated by dark purple arrowheads, nuclear factors are indicated by light purple arrowheads, chromatin is indicated by gold arrowheads, NE and NPCs and NE are annotated accordingly. Scale bar = 50 nm.

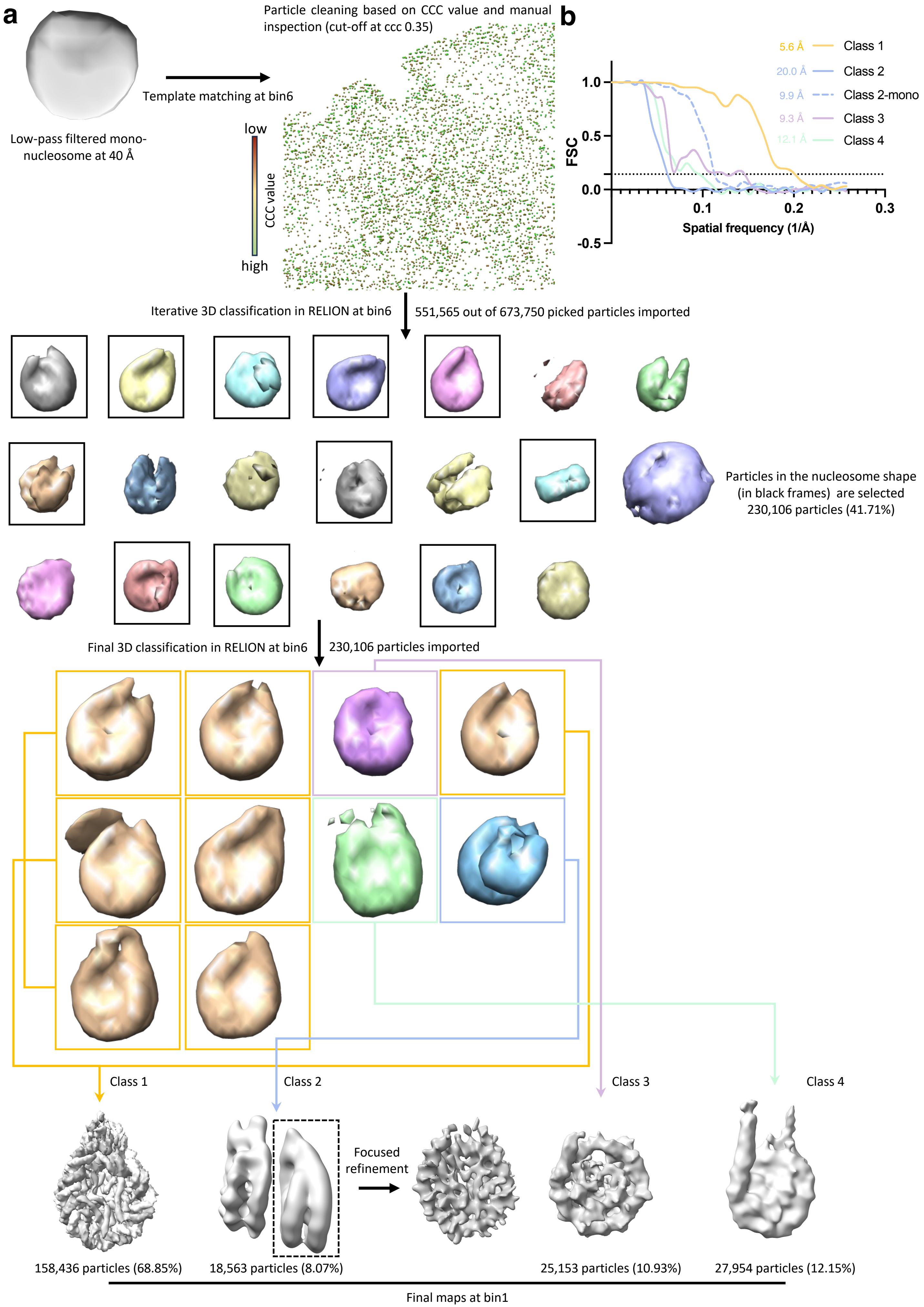

a
Low-pass filtered mono-nucleosome at 40 Å
Template matching at bin6
Particle cleaning based on CCC value and manual inspection (cut-off at ccc 0.35)
low
CCC value
high
b
FSC
Spatial frequency (1/Å)
1.0
0.5
0.0
-0.5
0.1
0.2
0.3
5.6 Å Class 1
20.0 Å Class 2
9.9 Å Class 2-mono
9.3 Å Class 3
12.1 Å Class 4
Iterative 3D classification in RELION at bin6
551,565 out of 673,750 picked particles imported
Particles in the nucleosome shape (in black frames) are selected
230,106 particles (41.71%)
Final 3D classification in RELION at bin6
230,106 particles imported
Class 1
Class 2
Class 3
Class 4
Focused refinement
158,436 particles (68.85%)
18,563 particles (8.07%)
25,153 particles (10.93%)
27,954 particles (12.15%)
Final maps at bin1

**Supplementary Figure 3 | Workflow of subtomogram averaging of nucleosomes**. **a**, The workflow used in this study for subtomogram averaging of nucleosomes. A low-pass filtered (40 Å) template is applied in the initial template matching using emClarity/1.5.0.2. The particle-picking number was intentionally set to an excessive value in selected regions (nuclear regions) of tomograms to ensure an exhausting search, yielding 673,750 particles in total. The matched particles were initially cleaned in Chimera thresholding the cross-correlation coefficient (ccc) value with a cut-off at 0.35, followed by manual inspection to further remove false positives residing outside the nucleus. The cleaned particles (551,565) are then transferred into RELION/4.0 for 3D classification using C1 symmetry, classes with a nucleosome shape (230,106 particles) are selected for the following iterative 3D classification until no obvious junk classes. **b**, Gold-standard Fourier shell correlation (FSC) curves of subtomogram averaged maps from nucleosome classes, H1-bound nucleosome (class1, gold), stacking H1-bound nucleosomes (class2, blue), H1-bound nucleosome in class2 (dashed blue), core nucleosome (class 3, purple), and open-linker H1-bound nucleosome (class 4, green). The resolution is indicated at 0.143 FSC cut-off.

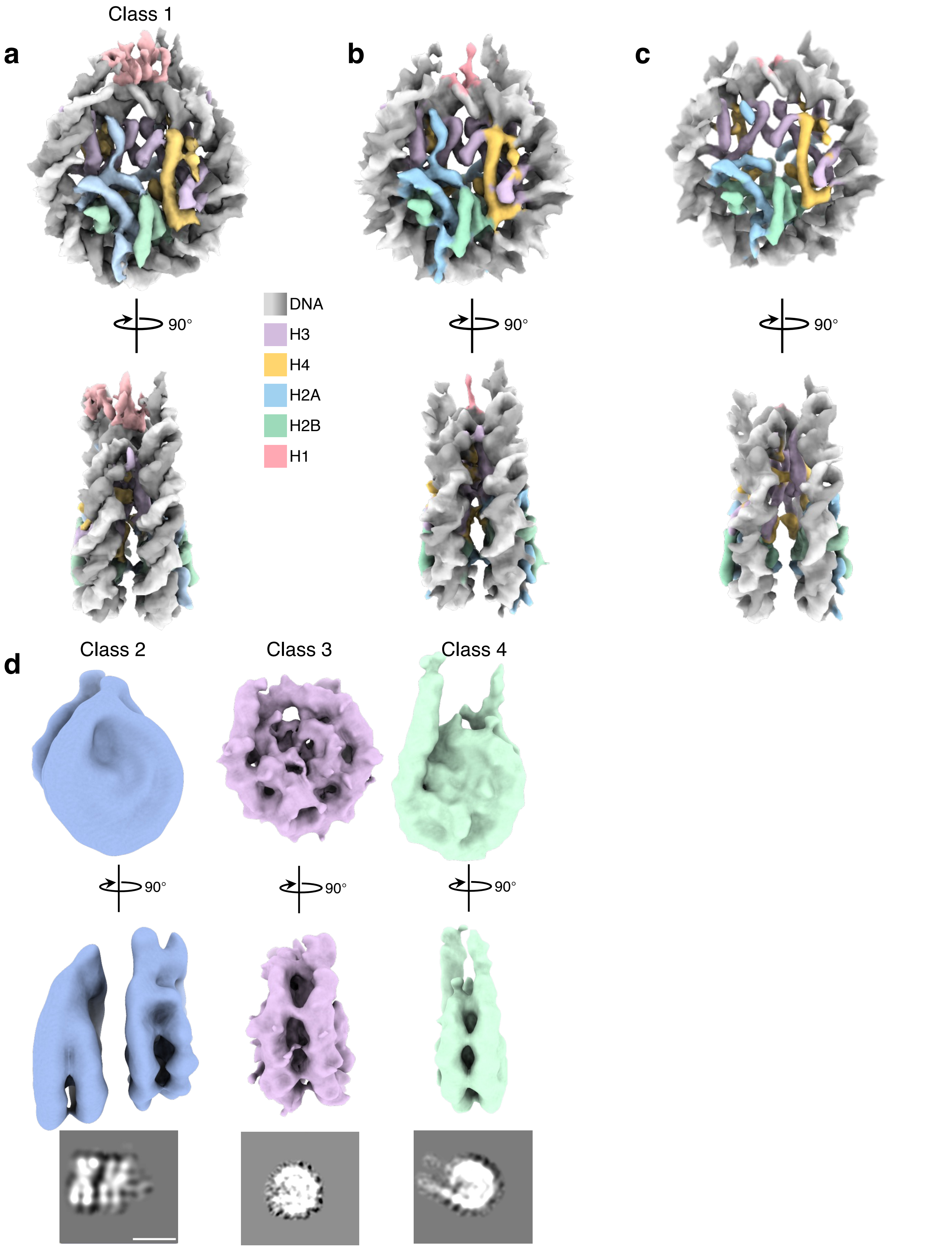


**Supplementary Figure 4 | In situ structure of nucleosomes. a-c**, In situ structure of the major nucleosome class (class 1) with histones and DNAs coloured: DNA (shades of grey), H3 (violet), H4 (gold), H2A (blue), H2B (green), and H1 (pink). Two orthogonal views are displayed with core histones shown at 6σ, and linker DNA shown at 5σ, and H1 shown at 4σ **(a)**, all contoured at 6σ **(b)** and all contoured at 8σ **(c)**. **d**, EM density maps of class 2, 3, and 4 contoured at the same level in **Figure 2c**. The bottom panel depicts central slices of EM maps.

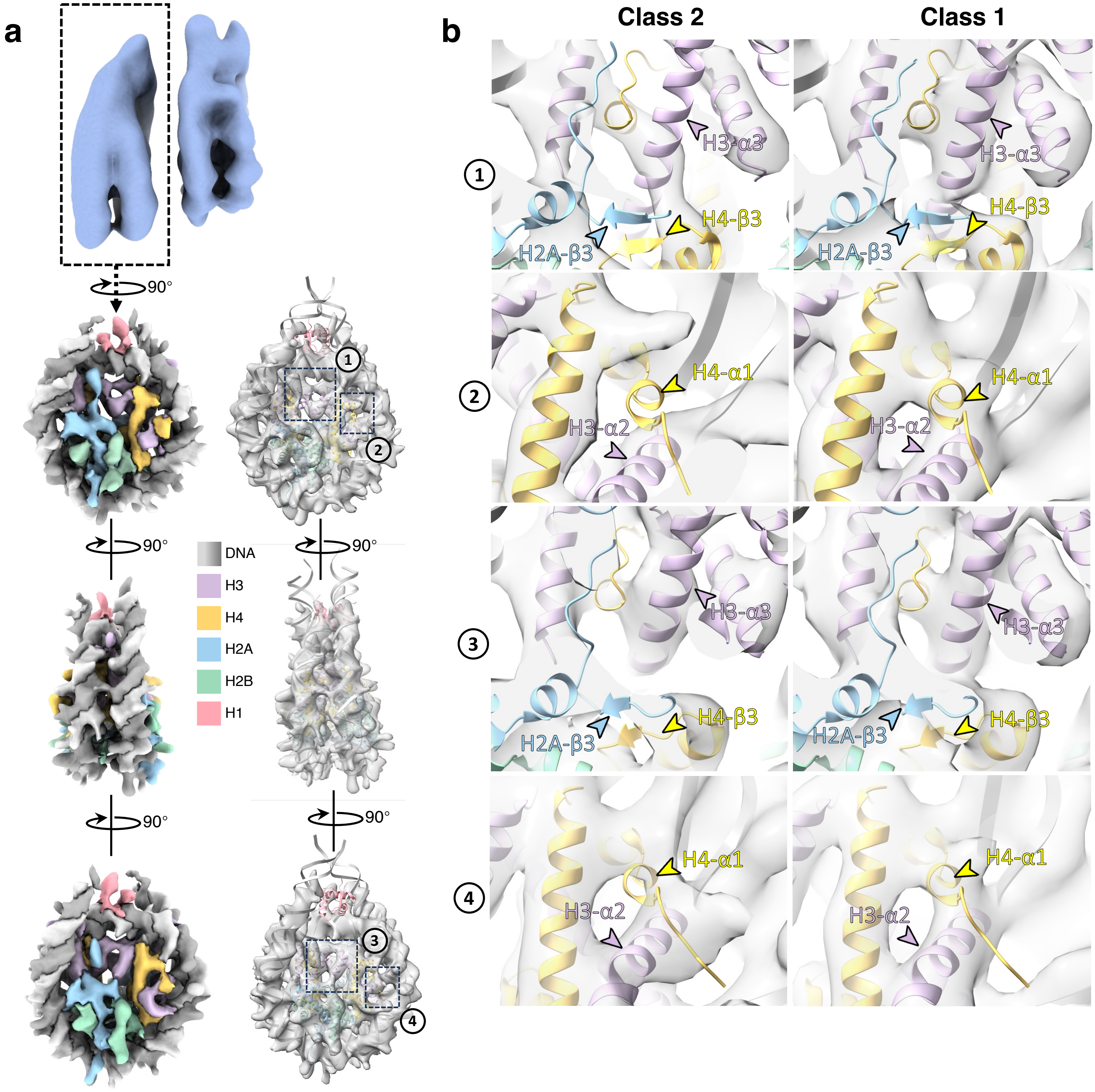


**Supplementary Figure 5 | Structural characterisation of stacking H1-bound nucleosomes. a**, In situ structure of the H1-bound nucleosome (framed by dashed box) in the stacking ones. Histones and DNAs are coloured as indicated. Three orthogonal views are displayed and contoured at 5.5σ, fitted with the atomic model (PDB 7PEX) on the right. Two major conformational changes in the H3 and H4 on the inter-nucleosome interface when compared to class 1 are indicated by the dashed frames and numbered 1 and 2. Region 3 and 4 are on the neighbor-free side. **b**, Zoom-in views of the regions of focus-refined H1-bound nucleosome in the stacking ones compared with mono H1-bound nucleosome (class 1). In region 1, the density of H2A-β3 and H4-β3 is missing and the conformation of H3-α3 is changed in class2. In region 2, the direction of H4 tail is changed in class2. In region 3 and zone 4, no prominent conformational difference is observed.

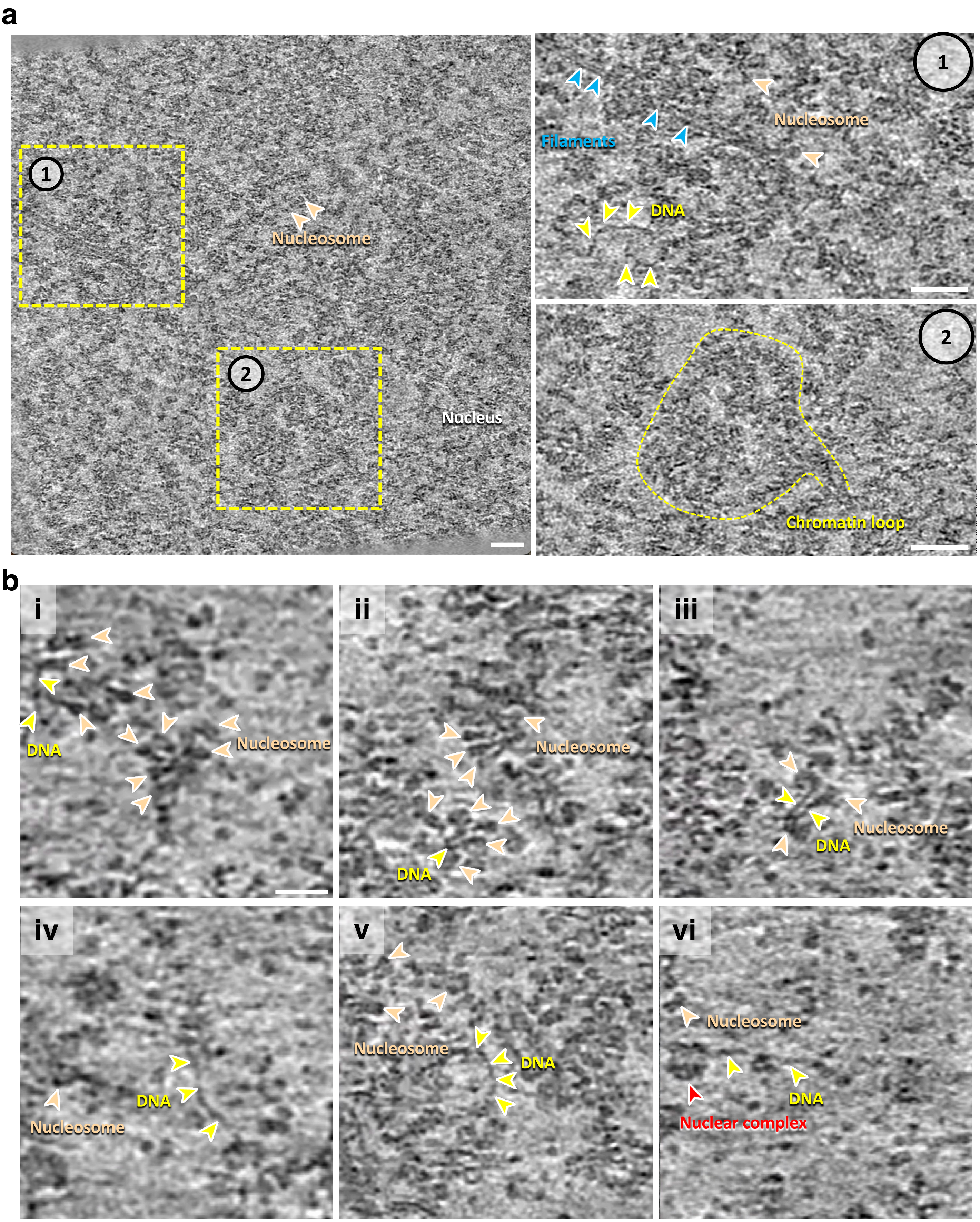


**Supplementary Figure 6 | Representative tomographic slices showing diverse structures of chromatin. a**, A representative tomographic slice of a tomogram deep in the nucleus without HIV-1. Two subregions are framed with prominent features: 1. Nuclear filaments connecting nucleosomes; 2. Chromatin loop. The two subregions are enlarged and displayed on the right. Scale bars = 50 nm. **b**, A gallery of diverse structures of chromatin. Nucleosomes are indicated by gold arrowheads, DNA is indicated by light yellow arrowheads, nuclear complex is indicated by the red arrowhead in **vi**. Scale bar = 20 nm.

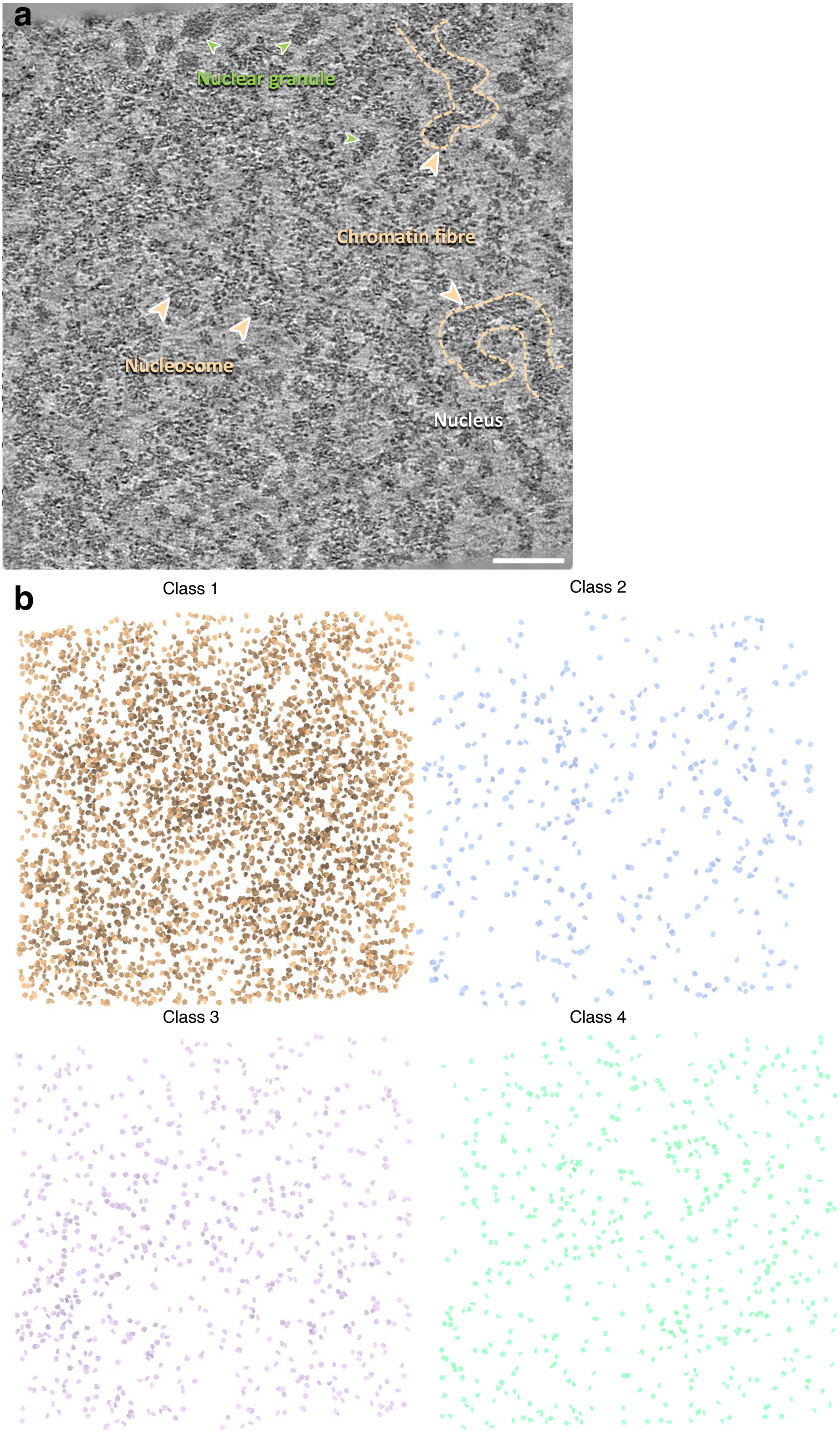


**Supplementary Figure 7 | Representative tomogram of chromatin without HIV-1. a**, A representative tomographic slice of a heterochromatin region in the nucleus. Nucleosomes and chromatin fibres are indicated by gold arrowheads, nuclear granules are labelled by green arrowheads. Scale bar = 100 nm. **b**, Map-back of individual classes in the heterochromatin region of the nucleus, with all classes included and coloured the same in **Figure 3d.**

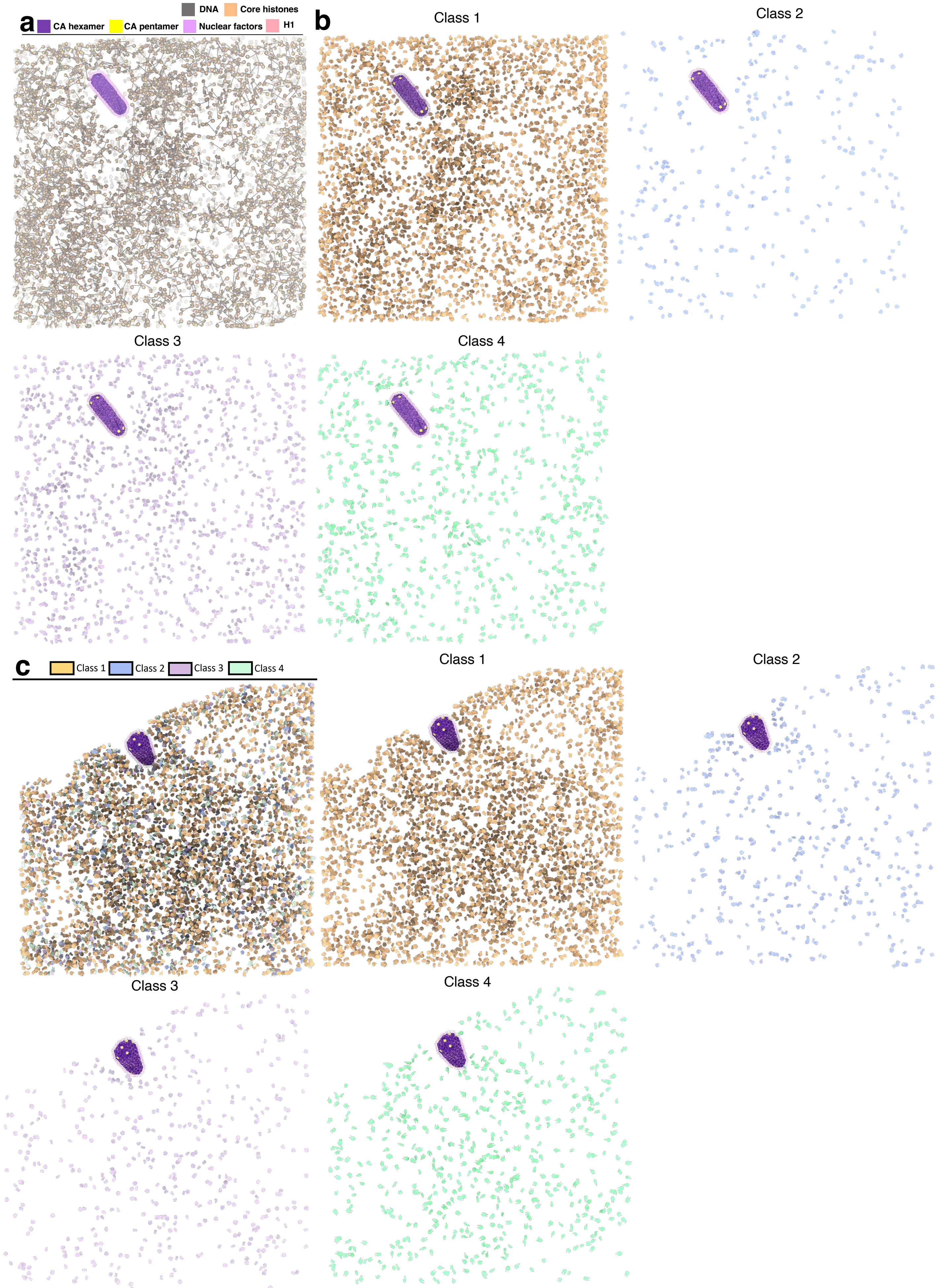


**Supplementary Figure 8 |Map-back of nucleosomes. a**, A map-back representation of nucleosomes linked by linear DNA strand of various lengths in a tomogram collected from the interior region of the nucleus containing a HIV-1 core, the same one in **Figure 1h**. Only zig-zag linked nucleosomes (see methods section) are highlighted for clarity. **b**, Map-back of individual classes in the interior region of the nucleus, with all classes included and coloured the same in **Figure 4c**. **c**, Map-back of nucleosomes by classes in the periphery region of the nucleus containing a HIV-1 core, the same one in **Figure 1d**, with all classes included and coloured accordingly.

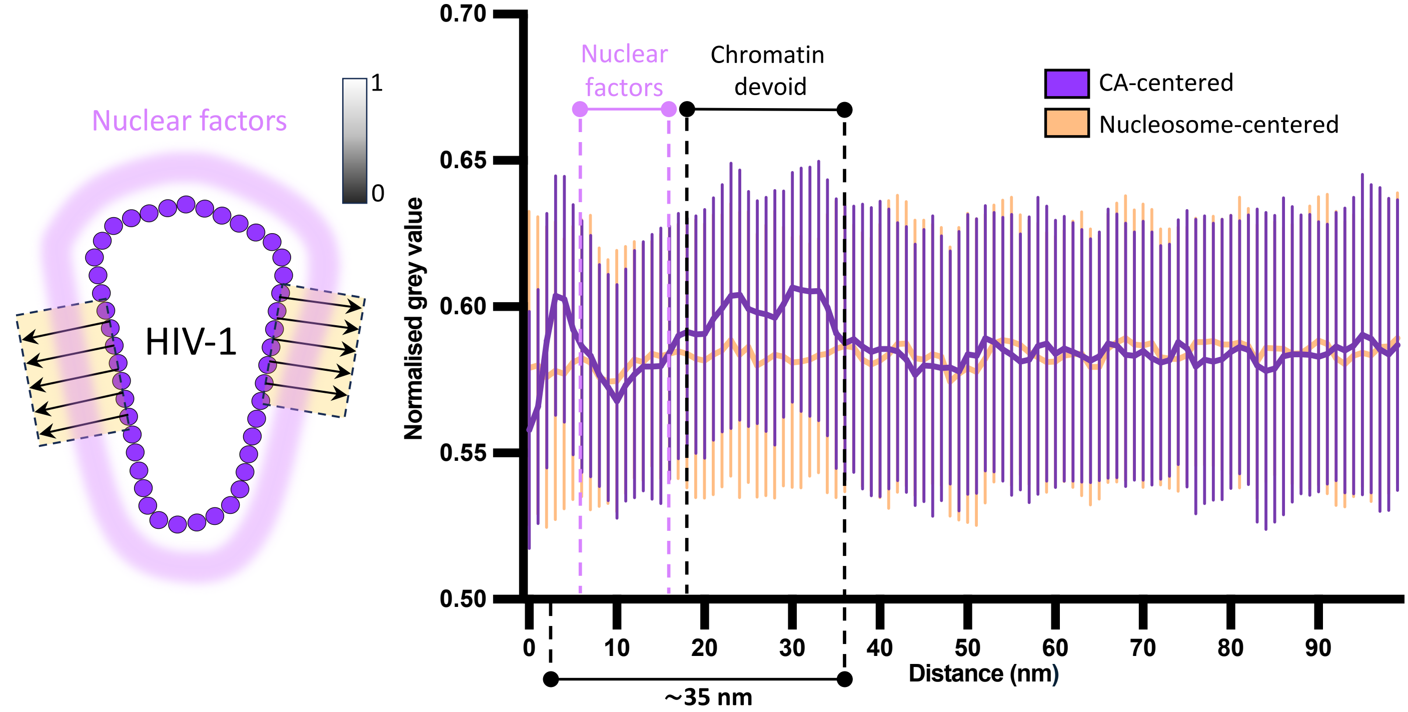


**Supplementary Figure 9 | Density profiling of the vicinity of imported WT HIV-1 cores.** A cartoon diagram (left) illustrates the measurement of grey values along the normal lines (arrows) extending from the core surface, including partial densities of CA (~3 nm). A control nucleosome centered density profiling plot is included. Lower intensity corresponds to higher density values. For WT cores, n = 233 lines × 12 cores. For control nucleosomes, n = 227 lines × 12 tomograms. The central lines represent the mean, and whisks represent the S.D.. Areas occupied by nuclear factors and devoid of chromatin are indicated accordingly.

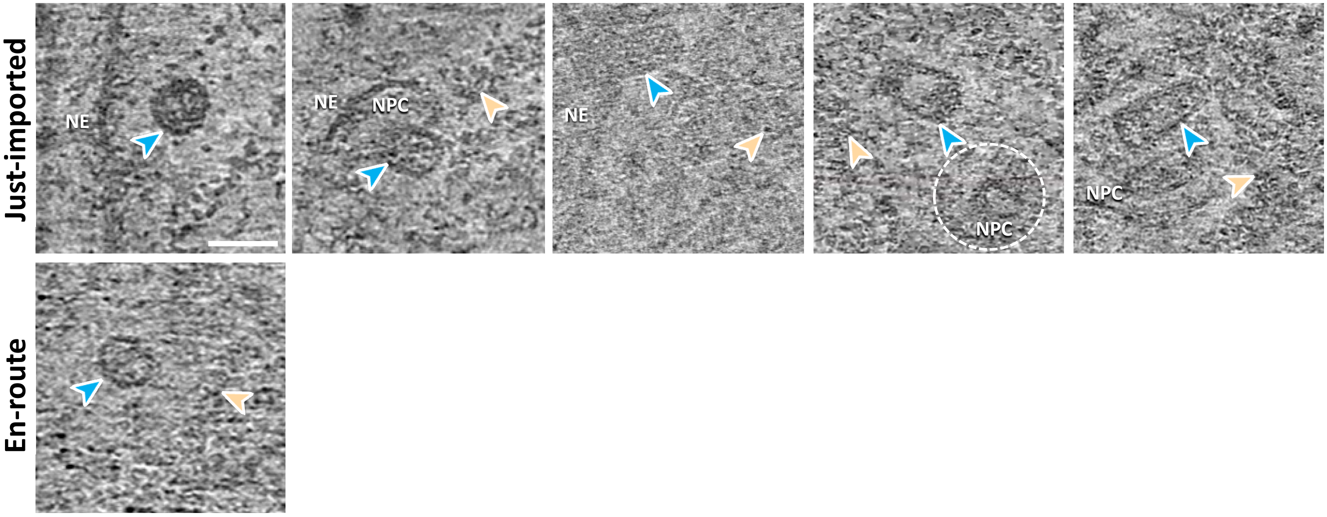


**Supplementary Figure 10 | Gallery of imported N74D HIV-1 cores.** N74D cores are indicated by light blue arrowheads, chromatin is indicated by gold arrowheads. NE and NPCs are annotated accordingly. Scale bar = 50 nm.

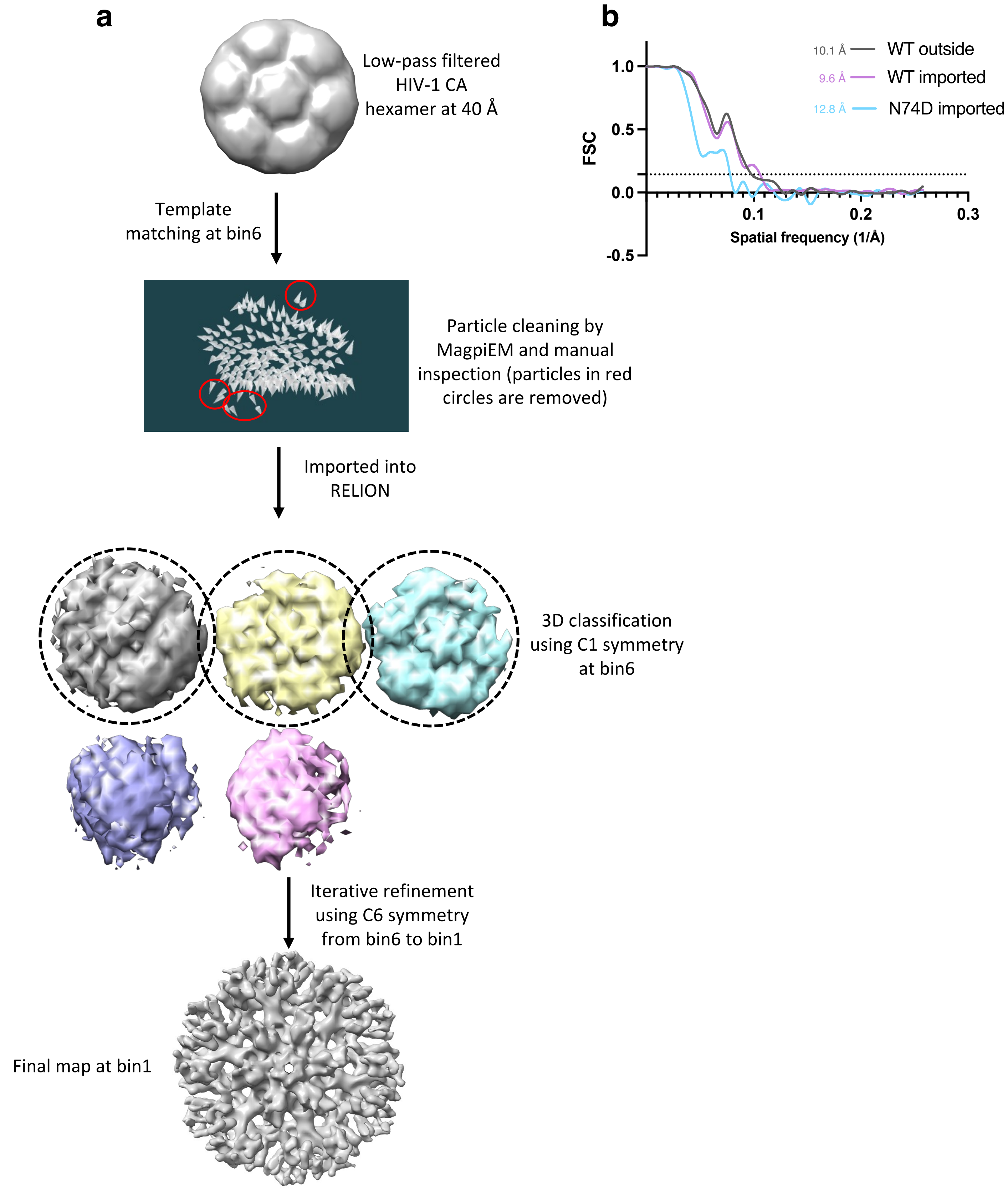


**Supplementary Figure 11 | Subtomogram averaging of HIV-1 CA hexamers**. **a**, The workflow for subtomogram averaging of CA hexamers. A low-pass filtered (40 Å) template is applied in the initial template matching using emClarity/1.5.0.2. The particle-picking number was intentionally set to an excessive value in cropped tomograms to ensure an exhausting search. The matched particles were initially cleaned using MagpiEM, followed by manual inspection in Chimera to further remove false positives (red framed particles). The cleaned particles are then transferred into RELION/4.0 for 3D classification using C1 symmetry, classes in a shape of CA hexamer are selected for the following iterative refinement until bin1. **b**, Gold-standard Fourier shell correlation (FSC) curves of subtomogram averaged maps from WT outside (grey), WT imported (light purple), and N74D imported (light blue) CA hexamers. The resolution is indicated at 0.143 FSC cut-off.

**Supplementary Table 1|** Cryo-FIB lamella preparation

| Microscope | Conventional cryo-FIB/SEM Aquilos 2 | Plasma cryo-FIB/SEM Arctis |
|---|---|---|
| **Voltage (keV)** | 30 | 30 |
| **Ion beam source** | Gallium | Argon |
| **Sputtering coating prior to milling (seconds)** | No | 30 |
| **GIS coating time (second)** | 30 | 50 |
| **Bulk milling current** | N/A | N/A |
| **Milling current** | 0.1-0.5 nA | 0.74-2 nA |
| **Polishing current** | 30 pA | 60 pA |
| **Sputtering coating post polishing (seconds)** | No | No |
| **Fluorescence microscope** | METEOR (50 ×) | iFLM (100 × ) |
| **Number of lamellae** | 30 | 15 |

**Supplementary Table 2|** Cryo-ET data collection

| Sample | Lamellae of WT HIV-1 cores | Lamellae of N74D cores |
|---|---|---|
| **Microscope** | FEI Titan Krios G3 | FEI Titan Krios G3 |
| **Voltage (keV)** | 300 | 300 |
| **Detector** | Falcon 4i | Falcon 4i |
| **Energy-filter** | Selectris X | Selectris X |
| **Slit width (eV)** | 10 | 10 |
| **Super-resolution mode** | No | No |
| **Physical pixel size (Å/pixel)** | 1.94 | 1.903 |
| **Defocus range (μm)** | -2 to -5, increment 0.25 | -2 to -5, increment 0.25 |
| **Acquisition scheme** | Dose-Symmetric, tilt span of 54$^{o}$, increment 2$^{o}$ step, group 3 | Dose-Symmetric, tilt span of 54$^{o}$, increment 2$^{o}$, group 3 |
| **Total dose (electrons/Å$^2$)** | 137.5 | 137.5 |
| **Number of frames** | 10 | 10 |
| **Number of lamellae** | 36 | 9 |
| **Number of tomograms** | 253 | 45 |

**Supplementary table 3|** STA of nucleosomes and HIV-1 CA hexamers

| Names | H1-bound nucleosome | Stacking H1-bound nucleosomes | H1-bound nucleosome in stacking ones | Core nucleosome | Open-linker H1-bound nucleosome | Outside WT CA | Imported WT CA | Imported N74D CA |
|---|---|---|---|---|---|---|---|---|
| **Particle number** | 158,436 | 18,563 | 18,653 | 25,153 | 27,954 | 6,404 | 4,445 | 988 |
| **Resolution by gold-standard FSC cut off (Å)** | 5.6 | 20.0 | 9.9 | 9.3 | 12.1 | 10.1 | 9.6 | 12.8 |
